# Characterizing Transient Protein-Protein Interactions by Trp-Cys Quenching and Computer Simulations

**DOI:** 10.1101/2022.08.05.502984

**Authors:** Lim Heo, Katukurunde Gamage, Gilberto Valdes-Garcia, Lisa J. Lapidus, Michael Feig

## Abstract

Transient protein-protein interactions occur frequently under the crowded conditions encountered in biological environments, yet they remain poorly understood. Here, tryptophan-cysteine quenching is introduced as an experimental approach that is ideally suited to characterize such interactions between proteins with minimal labeling due to its sensitivity to nano- to microsecond dynamics on sub-nanometer length scales. The experiments are paired with computational modeling at different resolutions including fully atomistic molecular dynamics simulations to provide interpretation of the experimental observables and add further insights at the molecular level. This approach is applied to model systems, villin variants and the drkN SH3 domain, in the presence of protein G crowders. It is demonstrated that Trp-Cys quenching experiments are able to not only distinguish between overall attractive and repulsive interactions between different proteins, but they can also discern variations in interaction preferences at different protein surface locations. The close integration between experiment and simulations also provides an opportunity to evaluate different molecular force fields for the simulation of concentrated protein solutions.

**Significance Statement:** Biological environments typically involve a variety of different proteins at very high concentrations where non-specific interactions are unavoidable. These interactions may go beyond simple crowding effects and involve transient contacts that may impact structure, dynamics, and ultimately function of proteins *in vivo*. While computer simulations have partially characterized such interactions, experimental data remain limited because established techniques are generally not well-suited to the characterization of dynamic processes on microsecond time and nanometer length scales. Tryptophan quenching by cysteine is introduced here as a new approach for studying transient protein encounters under concentrated conditions with the support of computational modeling. The study demonstrates that such experiments can resolve not just differences between different proteins but also residue-specific interaction preferences.

## Introduction

Biological cells are densely packed with proteins and other biomolecules at typical volume fractions of 20-40% in the aqueous phase (1). Frequent encounters between diffusing proteins are an unavoidable consequence at such high concentrations, but the full implications for biological function remain unclear. More specifically, it is necessary to understand how exactly protein diffusion, structure, and dynamics is affected at high concentrations (2–7). Another important dimension is that interactions at high concentrations may lead to aggregation and phase separation (8, 9).

A starting point for understanding crowding effects is the excluded-volume effect where diffusion is limited and where compact arrangements are favored for the lack of free space (10). Beyond this simplified scenario, recent studies have provided evidence that proteins may interact transiently to different degrees as a function of protein and concentration (11–15). Supported also by computer simulations (16–19), the idea is that proteins form dynamic clusters that persist on nanosecond to microsecond time scales, long enough to affect diffusion properties and interfere with ligand binding events (20). In fact, the argument has been made that the slow-down in diffusion experienced by proteins in crowded environments is primarily due to slower-diffusing clusters formed transiently and much less due to increased solvent viscosity or reduced free space (19, 21). Experimental evidence has come primarily from analyzing diffusion in concentrated protein solutions, with the key observations that diffusion varies as a function of protein crowder species at the same volume fractions (22–24), and that rotational diffusion is slowed down as much as or more than translational diffusion (12), a hallmark of interacting particles. Other approaches have explored transient interactions more directly, especially for specific biological systems (15, 25–28). It is still largely unclear, though, what exactly determines how strongly and in what manner two arbitrary proteins interact under highly concentrated conditions.

A direct characterization of the transient interactions between any pair of proteins via experiment has been difficult as the interactions are expected to be short-lived and presumed to lack strong preferences for specific intermolecular contact interfaces. In this work, we introduce quenching of the tryptophan (Trp) triplet state by cysteine (Cys) as a new approach for examining such interactions in concentrated solutions. Trp-Cys quenching has been used for about 20 years for investigating intramolecular dynamics of disordered polypeptide chains (29–34). The basic schematic of the technique is shown in **Fig. 1**. A Trp residue within a folded protein is excited to a long-lived triplet state via the fluorescence singlet state. In the absence of quenchers, the lifetime of this state is ~40 μs (30), but it is efficiently quenched via electron transfer only by Cys and not by other amino acids. Trp-Cys quenching is sensitive to microsecond dynamics over distances up to 10 Å, which is much shorter than characteristic distances of 20-70 Å in Fluorescence Resonance Energy Transfer (FRET) (35). At high physiological volume fractions, the most probable bimolecular distances are on the order of 10 Å (5), corresponding to about 1-3 water layers separating macromolecules (36), and closer when in contact. A FRET-labeled protein mixture would see almost 100% transfer with such short distances, and there would be little distinction whether molecules are in direct contact or not. In contrast, the Trp triplet in a concentrated solution may have a shorter lifetime than in dilute solutions, but it is still measurable, and one should be able to distinguish contacts from close interactions. Trp-Cys quenching is thus well-suited for characterizing transient intermolecular encounters in concentrated solutions. A further advantage is that only natural amino acids are involved and additional labeling that may interfere with protein-protein interactions can be avoided. Moreover, because of the sensitivity to relatively short distances, the location of Trp and Cys residues at specific locations on the protein surfaces provides spatial resolution that can be explored by moving Trp or Cys residues to other parts of a protein structure.

**Figure 1.**
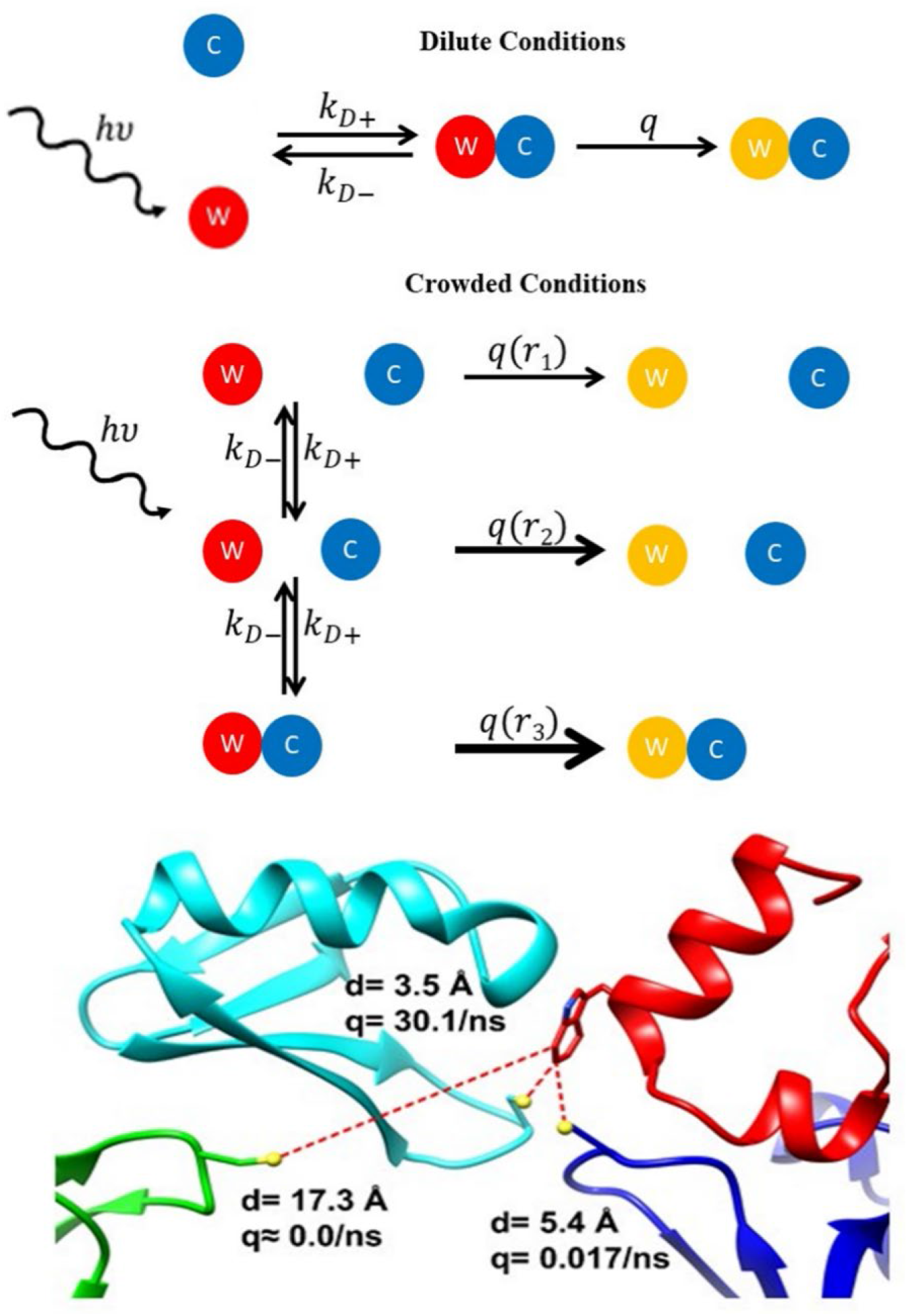
(top) Schematic representation of Trp triplet quenching by Cys. A protein containing Trp (orange ball) is excited to a long-lived triplet state (red ball) by absorption of a UV photon. If the Trp comes into contact with a Cys-containing protein (blue) ball, the excited state is quenched. Under dilute conditions, molecules are typically far enough away that after excitation of the Trp triplet state the Trp and Cys-containing proteins must diffuse towards each other before Trp is quenched to the ground state (yellow ball). Under crowded conditions, excitation of the Trp triplet state occurs when the Cys is already quite close, and quenching can happen at any point, but the rate will depend exponentially on distance between the proteins. (bottom) A snapshot of villin (red) in nearby contact with 3 protein G molecules (blue, green and cyan) is shown from the simulations with the Trp-Cys distances and quenching rates indicated.

As Trp-Cys quenching decays in themselves are difficult to interpret at the molecular level, computer simulations are needed to provide context. Molecular dynamics simulations of concentrated protein solutions in full atomistic detail and over microsecond time scales are possible now (37, 38), and this work presents a close comparison between experiment and simulation, including an opportunity for validating the force field parameters that are being used in the simulations. Once validated, the simulations can reveal additional information about protein-protein interactions at the molecular level. Together, this integrated experimental-computational approach provides direct, experiment-driven insights into transient protein interactions at elevated protein concentrations.

We focus here on well-known model systems, *i.e*. the chicken villin headpiece and the drkN SH3 domain as probe proteins with the B1 domain of protein G as the quencher protein that is present at high concentrations. The relatively small systems facilitate extensive computer simulations, but the approach described here is not limited in principle to such systems and can be applied to a wide variety of systems involving dynamic protein-protein interactions on short time scales and at close distances.

## Results

### Trp-Cys quenching experiments

Trp-Cys quenching experiments were carried out for villin or SH3 probe molecules in the presence of protein G quenchers. **Fig. 2A** shows the decay curves for the K33W variant of villin at different concentrations of protein G along with the derivative of the absorbance vs. log(*t*). In the absence of protein G (0 mM concentration), the decay curve reflects the unquenched lifetime of the Trp triplet state (40 μs). The slightly non-exponential decay is due to the presence of low-probability quenching by other amino acids within the folded structure. Once protein G is added, decay occurs on shorter time scales due to quenching. The resulting quenching rates in the presence of protein G clearly display non-exponential behavior. On a plot of absorbance vs. log(*t*), exponential kinetics manifests itself as constant absorbance at short and long times but with a sharp decay at the lifetime. The derivative of such a plot will have a sharp dip at the lifetime and will be zero at all other times. For a non-exponential decay, the dip in the derivative will be broader and shallower. Subsequent analysis of the quenching curves will focus on the depth and time location of the dip in the derivative.

**Figure 2.**
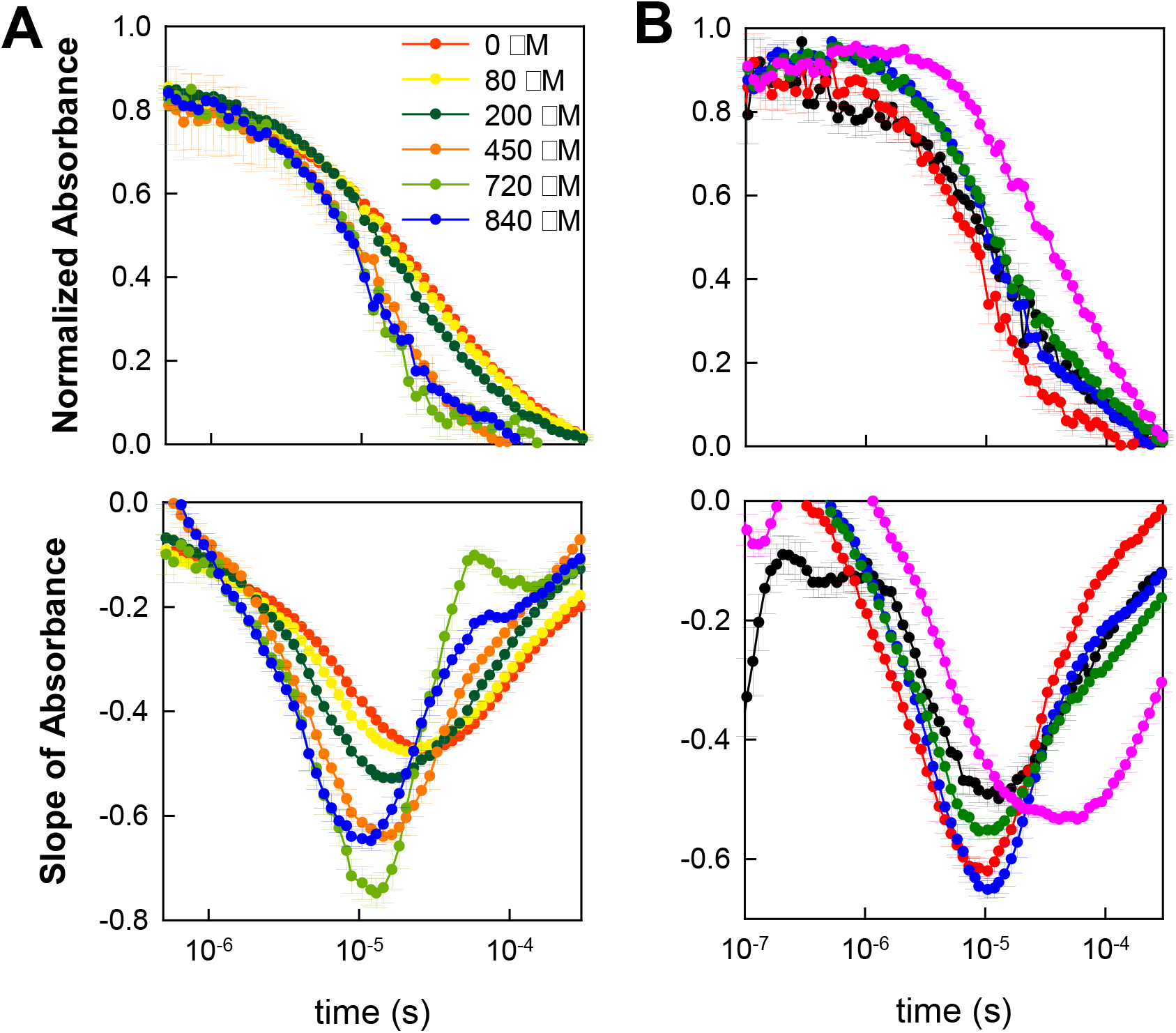
Experimental measurements of Trp triplet lifetime in villin variants and SH3 in the presence of protein G. Decays are shown as absorbance vs. time (top) and as the derivatives of the decays with respect to log(*t*) calculated as the slope of a line fit over a sliding window of 21 time points (bottom). (A) K33W at various concentrations of protein G. (B) Comparison of villin headpiece wild type (black), V10W (red), K33W (blue), R15T+K30E (green) and SH3 (magenta). Decay curves were normalized to 0.88 at 147 ns and to 0 at 369 μs. Each trace is an average of 6 independent measurements and the error bars represent the standard deviation. Error bars in the derivatives represent the error of the fit.

When protein G concentration is increased (**Fig. 2A)** both the location of the dip in time and the depth are affected. The minimum is shifted to shorter times at higher concentrations, because increased concentrations increase the frequency of protein-protein contacts. The depth changes as quenching shifts from slow, but short-range intramolecular quenching by other amino acids in the villin to faster intermolecular quenching by cysteine from interacting protein G molecules as shown for the mutant K33W in **Fig. 2A**.

In contrast, decay curves for different villin mutants at a fixed protein G concentration have varying depths of the derivative dip (**Fig. 2B** and **Table S1)** but with little shift in time. As will be shown more clearly below, the depth of dip is characteristic of the close-range interactions between the probe and quencher. This suggests differences in local interactions with respect to the Trp location (residue 24 in wildtype villin and the R15T+K30E mutant, and residues 10 and 33 in the V10W and K33W mutants, respectively), but overall similar association kinetics between protein G and the villin variants. On the other hand, the quenching curve for the SH3 domain is shifted significantly to longer times, but with a derivative minimum similar to the W24 villin mutants (**Fig. 2B** and **Table S1**), indicating slower SH3-protein G association kinetics compared to villin.

To check that proteins remain folded, we carried out CD spectroscopy and additional simulations. The main finding is that although some fraction of partially unfolded states may have been present, they do not appear to significantly affect the quenching results reported here (see Supplementary Material).

### Modeling of Trp-Cys quenching on 1D potentials

To better understand the determinants of the experimental quenching curves, we begin by studying a simple computer model where we can manipulate contact probabilities and association kinetics for probe-quencher pairs. The model involves a one-dimensional variable, the probe-quencher distance, that is sampled stochastically via Monte Carlo simulation for a given potential. **Fig. 3B** shows the resulting quenching curves obtained with a potential (**Fig. 3A**) that was matched to reproduce Trp-Cys distance distributions in the atomistic simulations described in more detail below. The shape and characteristics, including clear non-exponential behavior are similar as in the experiments. The curves are shifted in time when the time step in the time series is scaled, but also when the system size is reduced (**Fig. 3A/B**). A reduced system size is a model for increased concentration and the changes in the curves are similar as seen in the experiments at different protein G concentrations (**Fig. 2A**). It should be noted that the faster quenching at higher concentration is partially offset by slower diffusion due to crowding at higher concentrations and a full interpretation of the experimental data would involve scaling of the time in addition to a smaller system size to reflect increased concentrations.

**Figure 3.**
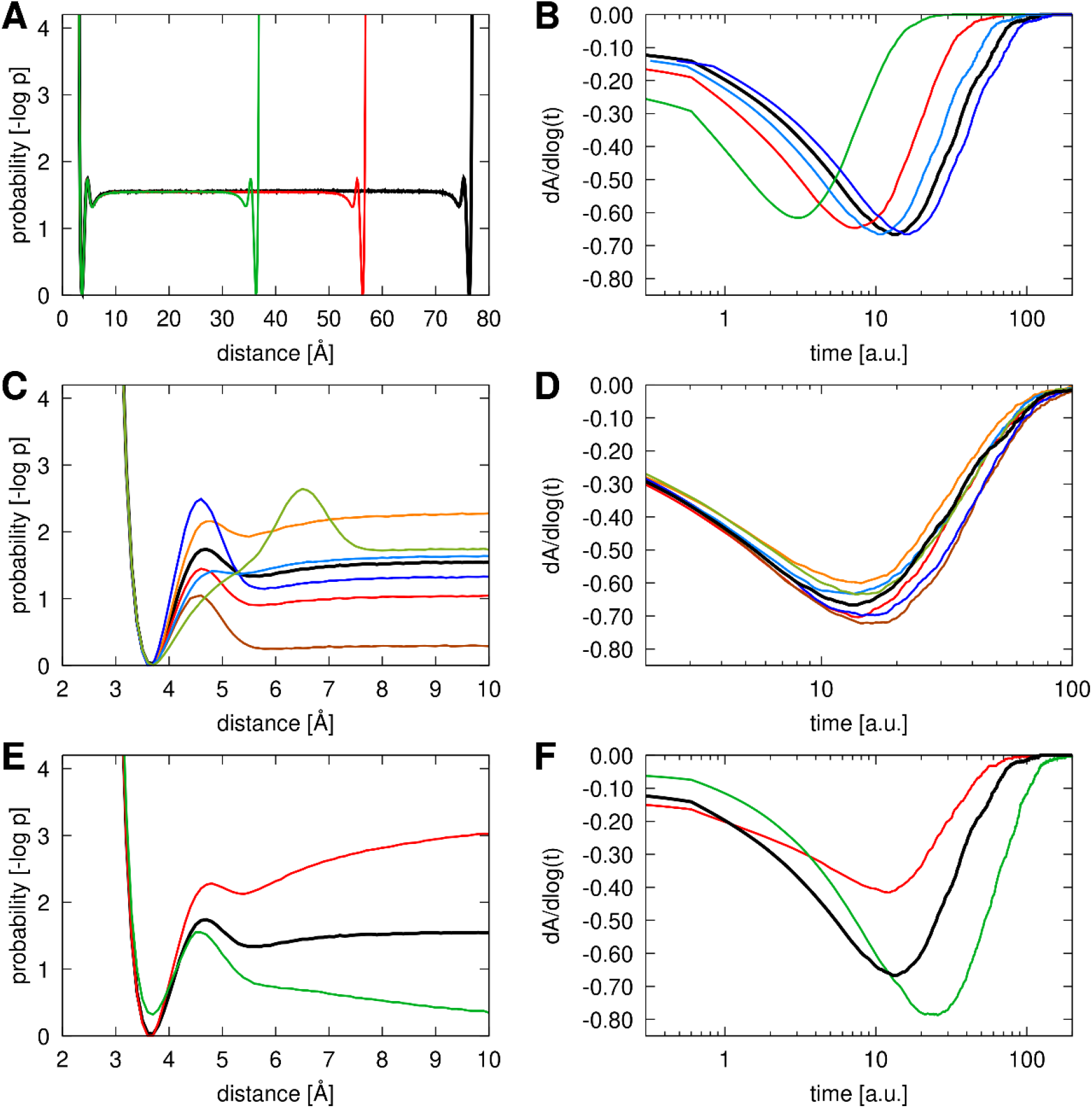
Computational modeling of Trp-Cys quenching via 1D potentials. Probability distribution functions are shown on the left (A, C, E) and derivatives of the resulting quenching curves are shown on the right (B, D, F). The reference distribution in black is similar to the contact probability extracted from atomistic simulations for the wild-type villin structure (see **Fig. 5**). Variations in system size (40 Å, black, 30 Å, red, and 20 Å, green) are shown in A and B. Panel B also shows additional quenching curves with scaled time (x1.2, light blue; x0.8, dark blue). Variations in the potential near the contact minimum are shown in C and D. Variations in long-range attraction (*d*=-1 in **Eq. S1**, red) or repulsion (*d*=1 in **Eq. S1**, green) are shown in E and F.

We focus the subsequent analysis on how the shape of the interaction potential affects the quenching curves. A variety of potentials were tested resulting in different contact minima depths and different kinetic barriers near the contact minimum. **Fig. 3C** shows examples of different potentials with a contact minimum and a barrier near the contact distance. The resulting theoretical quenching curves are shown in **Fig. 3D**. The curves shift little in time, and the main effect is a variation in the depth of the minimum in the derivative of absorbance vs. log(*t*). The depth is most strongly correlated with the difference in free energy between the contact minimum and large separation distances (calculated at 10 Å, **Fig. S1A**). A shallower contact minimum results in a deeper derivative value, a deeper contact minimum corresponds to a shallower derivative. The barrier height near the contact minimum, measured as the difference in energy at the peak vs. the energy at 10 Å, also has an effect as higher barriers shift the minimum down as well (see also **Fig. S1B**). In addition, smaller barriers shift the time slightly to shorter times whereas higher barriers slightly shift to longer times (**Fig. S1D**). This may be expected since potential barriers affect the association kinetics. If the potential is further modified by introducing long-range attraction or repulsion (**Fig. 3E**), the resulting quenching curves shift to longer time scales with a deeper minimum in the case of repulsion, but with long-range attraction the derivative mainly shifts upward (**Fig. 3F**). The shift to longer time scales in the case of repulsion is a result of less frequent contacts, whereas additional attraction does not significantly shift the time scale because most of the time is already spent in or near contact.

This analysis illustrates that the degree of non-exponentiality, quantified by the depth of the minimum in the derivative of the quenching curves, reports on the shape of the short-range interaction potential. The overall time scale of the decay curves reports generally on the dynamic time scale on which Trp-Cys interactions vary, but specifically captures longer-range features of probe-quencher interactions, meaning here beyond 10 Å. Broadly speaking, for different systems present at the same concentrations, a deeper minimum in the derivative corresponds to stronger local interactions at or near contact distances. Moreover, a shift to longer time scales indicates long-range repulsion.

This allows a first interpretation of the experimental data. The largest effect is seen between the SH3 domain and the villin variants where the quenching derivative curve for SH3 is shifted significantly to longer time scales. Following the above analysis, the data indicates long-range repulsion between SH3 and protein G relative to villin and protein G. This may be expected based on the overall electrostatics since both SH3 and protein G are negatively charged, and therefore repulsive, while villin and protein G have opposite net charges and are expected to be more attractive.

The more subtle variations between villin variants point to differences in interactions at specific locations of the Trp, *i.e*. near villin residue 24 (wild-type and R15T+K30E), near residue 10 (V10W mutant), or near residue 33 (K33W mutant). More specifically, it appears that protein G, or at least the surface patch near residue 10, where the Cys quencher is located, interacts less strongly with villin near villin residues 10 and 33 vs. 24 since the derivative minima are deeper for the V10W and K33W single mutants than for the wild-type (where the Trp is at residue 24). There is also a difference between the wild-type and double mutant (both of which have Trp at residue 24) indicating that the double mutant interacts less strongly. The double mutant reduces the positive charge of villin and therefore interactions with the oppositely charged protein G may be weakened, however, since the time scale in the quenching curves is not shifted between the two variants, we assume that the effect is more local instead of the long-range repulsion as proposed for SH3-protein G interactions.

### Atomistic simulations of Trp-Cys quenching

To interpret the experimental decay curves at the molecular level we applied μs-scale molecular dynamics (MD) simulations. The simulations sample the diffusion and interactions between the proteins in the same systems that were studied experimentally, *i.e*. villin mutants and the SH3 domain in the presence of protein G crowders. The survival probabilities of the triplet state of Trp were calculated from the simulation according to **Eq. 1** based on the distance distributions between Trp and Cys residues with the time-scale corrections described in the Methods section. They are compared with the experimental results in **Fig. 4**. For wild-type villin, we considered four CHARMM force field variants c36 (39), c36+water (16), c36m (40), and c36mw (40) (**Fig. 4A**). All force fields result in somewhat faster decay curves compared to experiment. That will be discussed in more detail below. Depending on the force field, the minimum depth in the derivative of the survival probability function varies. The deepest minimum is found for c36 with water scaling (c36+water) where the scaling increases protein-water interactions over the standard c36 force field (16), followed by c36mw (40), which also has increased protein-water interactions. Shallower minima are found for c36m (40) and c36 (39) (**Fig. 4A**). The analysis based on the 1D model above suggests that deeper minima indicate weaker protein-protein interactions. As expected, that is indeed the case as villin-protein G interactions are decreased with the c36+water and c36mw force fields compared to c36 and c36m according to protein-protein radial distribution functions (**Fig. S2**). The reduced interactions also result in smaller and shorter-lived clusters due to transient interactions (**Fig. S3**). Average cluster sizes that included villin WT in the cluster were 3.69, 3.21, 2.06, and 2.00 for c36, c36m, c36+water, and c36mw, respectively. c36 and c36m perform similarly, but the agreement with the experimental data is somewhat better for c36m. c36m has also been found to perform well in other contexts (40–42). Therefore, we will focus on simulations using c36m in the subsequent analysis.

**Figure 4.**
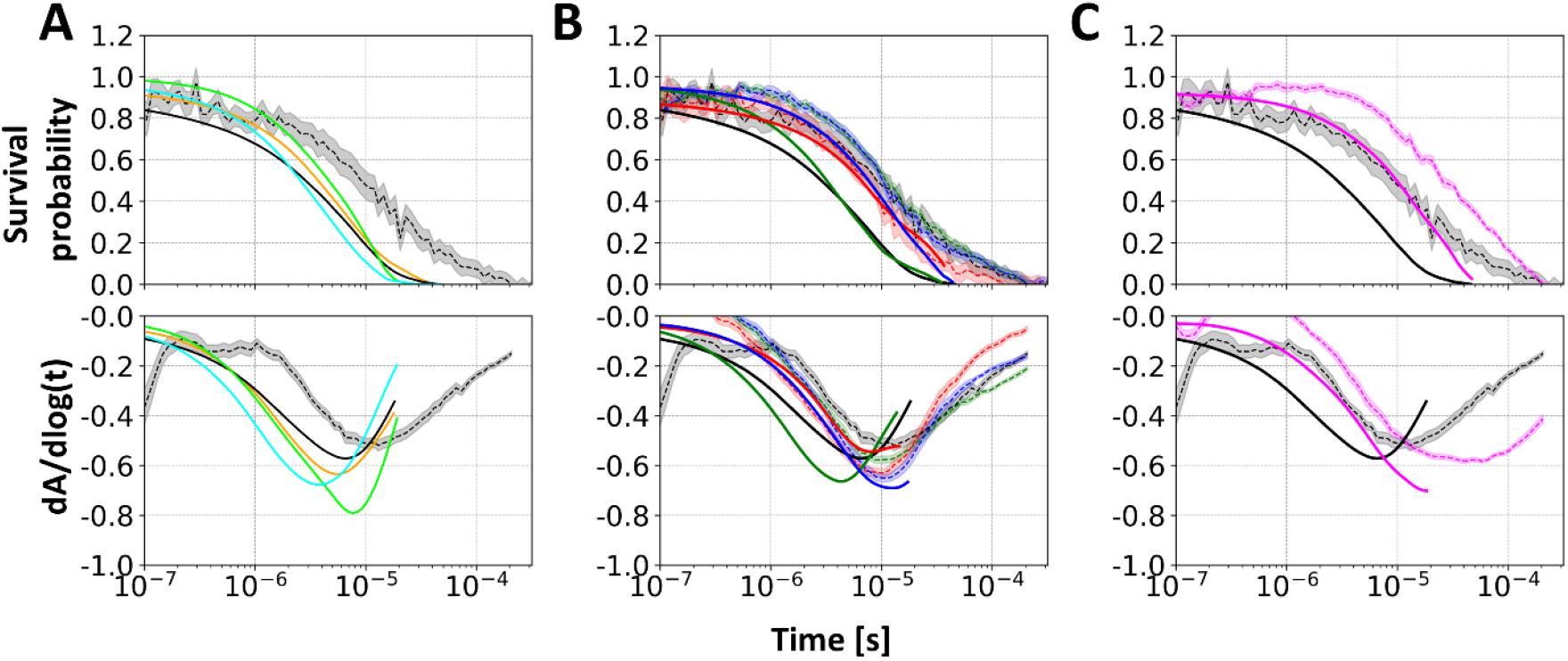
Comparison of survival probability decays between experimental measurements and simulated data. The probability and its derivative against log-time are shown at top and bottom of each panel, respectively. Derivative values were obtained from a linear-fit of probabilities against log-time with a window size of 21. Experimental values shown as solid lines with transparent shades for standard errors were adjusted to match the simulated data at 100-500 ns since the initial decay varies in the simulations and experiments are not sensitive to quenching during the initial 100 ns after excitation. A: Results for wild-type villin obtained using different force fields c36 (orange), c36+water (lime), c36m (black), and c36mw (cyan); B: Results for wild-type villin (black), V10W (red), K33W (blue), and the R15T+K30E double mutant (green); C: Results for wild-type villin (black) and SH3 (magenta).

Between different villin mutants, the simulations capture many of the features seen in the experimental quenching curves (**Fig. 4B**). The double mutant, R15T+K30E, has a deeper minimum in the derivative than the wild-type, indicating weaker local interactions that may stem from the less positive charge in the mutant (−1 vs. +2 for wild-type villin). Moreover, the K33W mutant shows weaker interactions than V10W as in the experiment (**Fig. 4B**). For both single mutants, the derivatives of the survival probabilities are in excellent agreement with experiment, but the minima of the wild-type and double mutant are both shifted downward compared to the experimental values. The time scale on which the triplet state survival probability decays is also matched for the single mutants but shifted to shorter times for the wild-type and double mutant. The shift in time scale for the wild-type and the double mutant (both of which probe Trp-Cys quenching with respect to Trp at villin residue 24) was unexpected and is not explained by differences in translational diffusion, as they are similar (**Table S2**). This will be revisited below.

The comparison between wild-type villin and SH3 based on the simulations shows a similar shift to longer times and with a deeper minimum (in SH3) as seen in experiment. SH3 is about 60% heavier than villin WT (molecular weights are 6811.5 and 4214.9 g/mol for SH3 and villin WT, respectively). The heavier mass of SH3 may suggest that it would diffuse slower, but in the simulations with crowders, both proteins diffused at a similar rate (**Table S2**), essentially because of less transient cluster formations between SH3 and protein G than between wild-type villin and protein G (**Fig. S4**). Again, differences in intermolecular diffusion do not explain the simulation results.

To further understand the simulation results, and ultimately the experimental data, we turn again to the 1D Monte Carlo model introduced above. **Figs. 5A/C** show the probability functions for Trp-Cys distances extracted from the atomistic simulations for the different systems studied here. In all cases, there is a clear contact minimum around 3.5-4 Å for villin, and somewhat broader for SH3, followed by a barrier at varying locations between 4.5 to 8 Å depending on the system, and a longer-range decline of the probability function. The decline reaches a plateau for villin mutants at about 15 Å but is significantly steeper for SH3. A depletion beyond 10 Å is generally expected as protein G favors interactions with villin due to crowding but may interact in many arrangements that do not place protein G’s Cys10 near the villin Trp. Beyond 25 Å, the probability function increases sharply since there is almost always a protein G molecule sufficiently close for minimum distances to be less than 30 Å due to crowding. Therefore, longer distances are not considered here.

**Figure 5.**
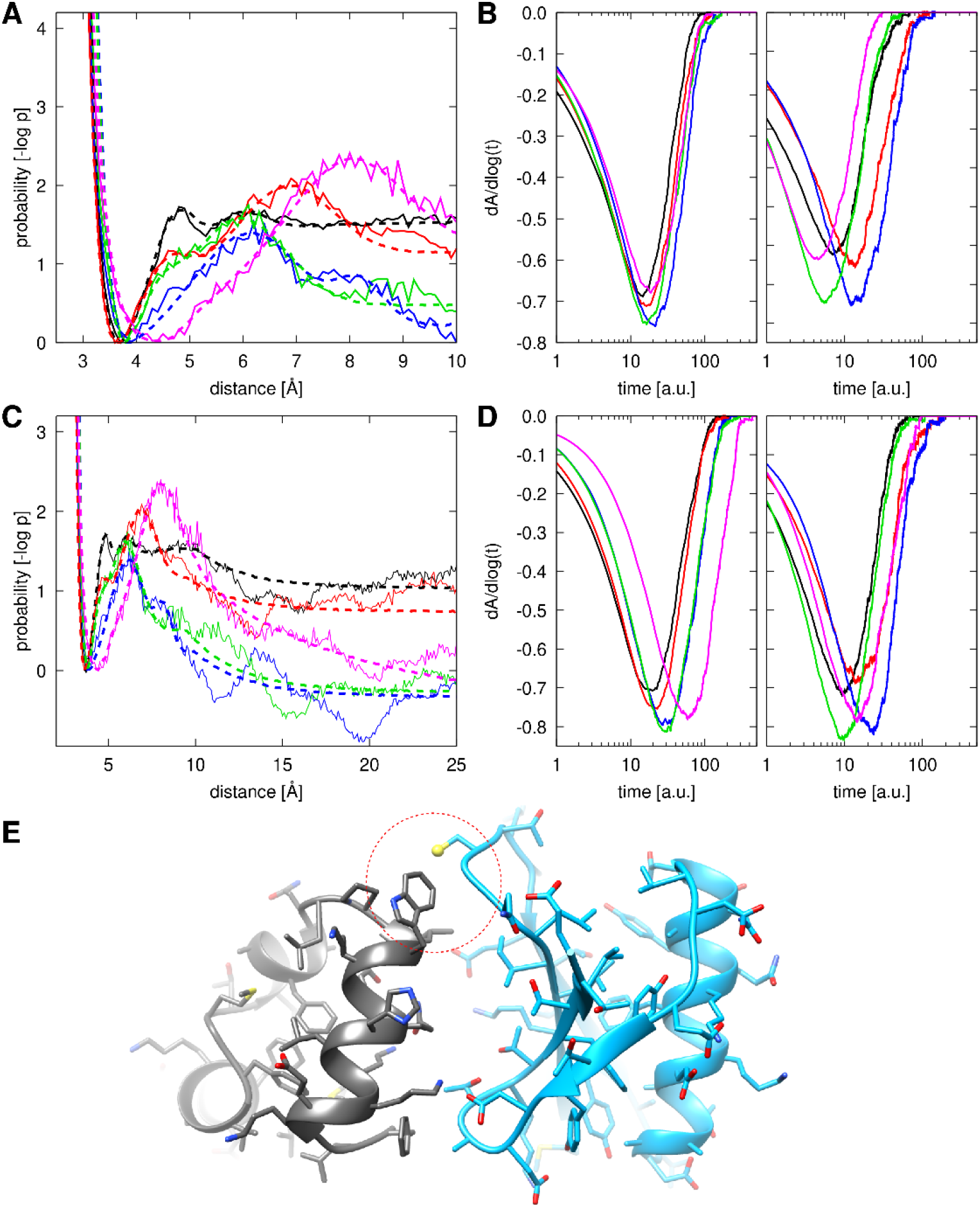
Trp-Cys contact probability functions (A, C) from atomistic simulations (solid lines) and fitted 1D potentials (dashed lines) and derivatives of quenching curves based on Monte Carlo sampling on 1D potentials (B, D). The atomistic probabilities are for minimum villin-protein G Trp-Cys distances. In the top panels (A, B), 1D potentials were fit to reproduce only the short-range behavior (<10 Å). In the center panels (C, D), potentials were fit to include the long-range behavior (up to 25 Å). Derivatives on the left were obtained with uniform Monte Carlo sampling, derivatives on the right resulted from sampling with diffusion-matched, distance-dependent step sizes (see text and Supplementary Material). Different colors reflect different probes: villin wild-type (black), villin V10W (red), villin K33W (blue), villin R15T+K30E (green), SH3 (magenta). The bottom panel (E) shows a typical interaction between protein G and wild-type villin when quenching occurs between Cys10 on protein G (blue) and Trp24 on villin (black).

1D potentials were fitted to the atomistic contact probabilities to obtain similar probability functions via Monte Carlo sampling (dashed lines in **Figs. 5A/C**, see details in the Supplementary Material). Quenching curves were then calculated from the Monte Carlo sampling (**Figs. 5B/D**). When only the short-range potential is considered (**Fig. 5B, left**), the time scales are similar for all systems and the derivative minima are ordered similar to experiment (*i.e*. the double mutant is lower than wild-type villin and V10W and K33W are lower than the double mutant). As discussed above, this is a direct consequence of the probability at close contact being more favorable compared to the probability at a distance of 10 Å (**Figs. 5A/C**) for the wild-type villin than the other villin variants. Once the long-range decay of the Trp-Cys contact probability is considered in the 1D model (**Fig. 5D, left**), the villin variants shift slightly in time, while maintaining their order, whereas the quenching curve for SH3 shifts more significantly to slower times and with a lower derivative minimum because of longer-range repulsion, qualitatively similar to the experimental observations.

So far, the Monte Carlo-sampled 1D potentials do not fully reproduce the quenching curves from atomistic simulations. In particular, the shift to shorter times seen for wild-type villin and the double mutant are not reproduced. To understand this observation, we analyzed diffusion in the Trp-Cys distance variable (**Fig. S5**). Diffusion in the Trp-Cys distance around 10 Å is about twice for wild-type villin and about five-fold for the double mutant and SH3 compared to diffusion in the V10W and K33W mutants at that distance (**Fig. S5**). In contrast, because of the Monte Carlo sampling protocol with a fixed maximum step size, the effective diffusion was not very different for the different systems (**Fig. S5B**). However, it is possible to impose different kinetics on the Monte Carlo sampling by adjusting the step size differently for different systems to reflect the variations seen in the atomistic simulations (cf. **Fig. S5D** and methodological details in the Supplementary Material). Once the 1D potentials were resampled with the altered pseudo-kinetics, the calculated quenching curves became much more similar to the results from the atomistic simulations (**Fig. 5B/D**, right) as wild-type villin, the double mutant, and SH3 were all shifted to shorter time scales relative to V10W and K33W. Interestingly, the depth of the minima also changed slightly, with the wild-type curves now lower than V10W as in the atomistic simulations indicating that different kinetics also has a small effect on the quenching curve derivative minima.

As mentioned already, there are only small differences in the translational diffusion values for the villin variants, SH3, and protein G and it is not immediately obvious why the Trp-Cys distance would fluctuate more rapidly with respect to Trp at the villin 24 position (probed by the wild-type and double mutant) vs. Trp at the 10 and 33 positions.

To understand this better, we first analyzed Trp-Cys contact lifetimes based on contact correlation analysis (**Fig. S6**). Generally, contacts persist on time scales of 5-30 ns, but the biggest difference is between SH3 and the double mutant and the other villin variants, likely indicating increased electrostatic charge repulsion but not explaining the difference in the quenching curves between wild-type villin and the V10W and K33W mutants. Overall cluster size distributions and lifetimes between wild-type villin and V10W/K33W are also similar and not explaining the clear difference in Trp-Cys distance diffusion (**Fig. S4**). Finally, we analyzed the conformational fluctuations of the Trp residue itself at different villin positions (**Fig. S7**). With or without protein G, Trp (or Tyr when substituted in V10W and K33W) is more dynamic at the 24 position than Trp at the 10 and 33 positions. The differences become significant at time scales beyond about 10 ns when protein G is present (**Fig. S4**). In the more concentrated systems studied via simulation, villin-protein G interactions persist for long enough times for differences in the Trp fluctuations as a function of where the Trp is located to matter. More specifically, we propose that the shift in the quenching rates for wild-type villin and the double mutant relative to V10W and K33W is a consequence of long associations between villin and protein G in arrangements as shown in **Fig. 5E** in which dynamics in the Trp-Cys distance is primarily due to fluctuations of the Trp residue. To further test this idea, we calculated hypothetical quenching curves relative to the residue at position 24 (Tyr for V10W and K33W) based on the V10W and K33W simulations. As shown in **Fig. S8**, these quenching curves are shifted also for the V10W and K33W simulations, similar to the wild-type and double mutants. These results show that it is the dynamics relative to the 24 position of villin that is determining the observed overall kinetics in the simulation instead of other differences due to the V10W and K33W mutations.

With a full understanding of how to relate the atomistic simulations to the experimental data collected at more dilute conditions, we finally turn to a more detailed interpretation of the experimental findings. While the reduced interactions between villin and SH3 and between villin and the double mutant are largely attributed to more repulsive electrostatic interactions, the differences between the wild-type and the V10W and K33W mutants are less obvious. **Fig. 6** shows that overall villin-protein G interaction preferences are similar for the three variants. However, interactions between protein G (more specifically, residue 10, where the Cys quencher is located) to villin residue 24 are apparently preferred over interactions near villin residue 10 and interactions near residue 33 (**Fig. 6F** and **Fig. S9**). This suggests that the Trp-Cys quenching experiments are an exquisitely sensitive tool to map out residue-specific interactions between probe-quencher protein pairs for transiently interacting proteins.

**Figure 6.**
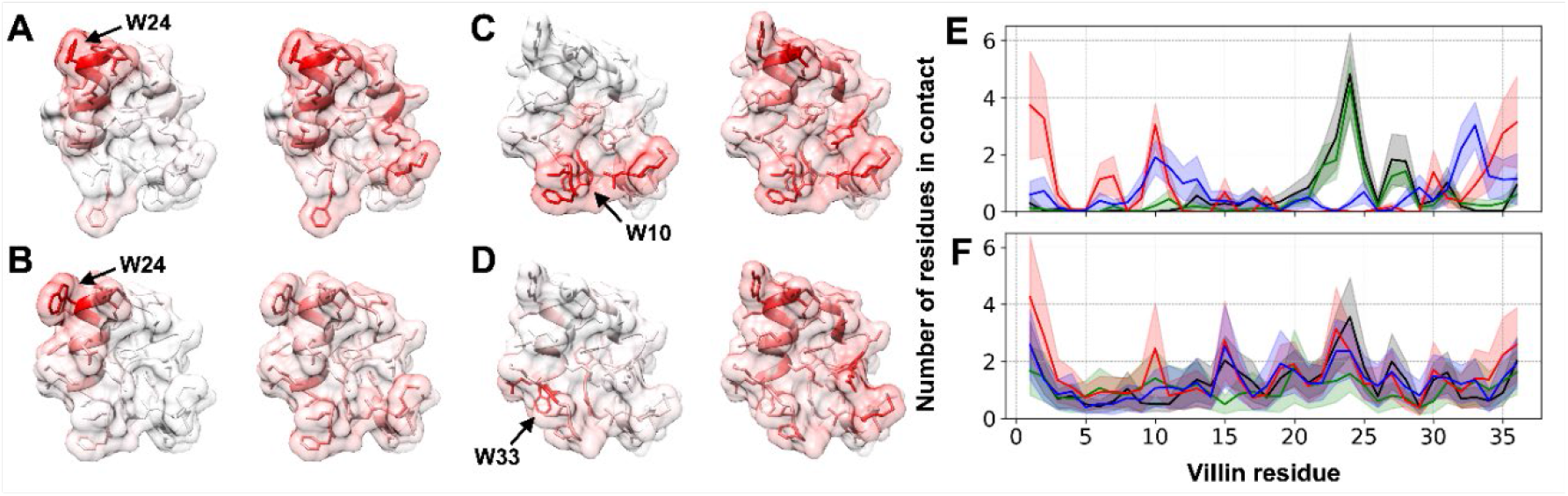
Residue-wise contacts between protein G and villin variants, projected onto the villin surface (A-D) and as a function of villin residue index (E, F). Surface projections are shown at the time of Trp-Cys quenching contact (left) and at any time of contact (right) for wild-type villin (A), the R15T+K30E mutant (B), the V10W mutant (C), and the K33W mutant (D). The location of the Trp residue is indicated by arrows. Contacts per frame vs. residue index are shown at the time of quenching contact (E) and at any time of contact (F) with different variants colored as in **Fig. 2**. Shaded areas indicate standard errors. Contacts were defined by residue pairs whose inter-atomic distances were closer than 5 Å.

Further analysis of protein G-villin contacts formed upon Trp-Cys quenching suggest that while quenching near residue 24 is highly specific to the immediate vicinity of residue 24 (**Fig. 6A/E**), quenching near V10W and K33W is more broad (**Fig. 6B,C,E** and **Fig. S10**) involving more residues as a result of the different location the residues on the villin structure on a flatter, broader surface compared to the more solvent-exposed 24 site at the tip of the villin structure.

From the perspective of the protein G quencher, there are fewer differences when interacting with different villin variants (**Fig. S11**). The vicinity of the Cys10 location is highly preferred for interactions with villin. Upon quenching, the residue interaction preferences become even more focused to this region of protein G. It does appear, though, that interactions are slightly more focused to the C10 residue on protein G when interacting with the 10 and 33 sites on villin compared to when protein G interacts with the wild-type/double mutant 24 quenching site (**Fig. S11**).

## Discussion

In this work, we have shown that Trp-Cys quenching can provide detailed information about transient protein-protein interactions. This technique has the unique features of sensitivity to close-range interactions and a strong distance dependence of the quenching rate, which illuminate the spatial distribution of proteins in a crowded environment. We show here that at fixed concentrations, we can use Trp-Cys quenching to map out interaction preferences between probe and quencher proteins with respect to different surface patches. Moving the location of the Trp to additional points on the villin surface could provide a more complete map of interaction preferences. Moreover, one could further probe interactions between villin and protein G by also moving the location of the Cys to other parts of the protein G surface. For example, one could move the Cys to a part of the protein G surfaces that is not predicted to interact extensively with villin. Such experiments should result in quenching curves with more exponential behavior, *i.e*. the derivative minima should be deeper without otherwise affecting the time scale of the quenching curves. This will be tested in future work. The comparison between villin and SH3 suggests that Trp-Cys quenching can distinguish between overall more or less attractive protein-protein interactions and further experiments could focus on a larger variety of protein pairs to map out interaction preferences.

The interpretation of the experimental data was initially guided by Monte Carlo modeling, showing that the overall shape of the quenching curve can be recapitulated with a small number of features in the potential of mean force over the first 10 Å of intramolecular distance. Ideally, one would like to turn this around and fit interaction potentials to given experimental curves. It may be possible to develop such a procedure in future work with certain assumptions about fixed concentrations and focusing on relative differences between different proteins in comparison to a known reference system.

In addition to the Monte Carlo modeling, all atom simulations allowed us to analyze the detailed intermolecular interactions between two proteins. The preference of protein G for villin near residue 24 was evident both experimentally, in the depth of the derivative curves, and computationally, in the probability of contacts and the simulations provided further insights into preferred interaction patterns between the probe and quencher proteins at the residue level.

Another important role of atomistic simulations is the prediction of dynamics of intermolecular interactions. The discrepancy between simulation and experiment for the W24 sequences of villin even after scaling for concentration shows that there are additional dynamic effects of crowding beyond the conventional “billiard ball” framework of diffusion. It appears that crowding-induced “sticky” interactions can significantly affect the dynamics of certain aspects of protein-protein interactions. This observation also highlights the need for experimental conditions to closely match the highly crowded conditions we wish to investigate. Protein solubility, especially at concentrations approaching biological cellular conditions, is a significant problem for *in vitro* experiments. This may be overcome in the future by creating condensates that induce millimolar concentrations at a nominal bulk concentration that is 10 times smaller. Our previous work on condensation of folded proteins and RNA shows that such condensates are possible (43) and may be employed in future work.

## Materials and Methods

### Systems

Sequence variations of the HP35 fragment of the villin headpiece (see **Table S3**) and the T22G mutant of the drkN SH3 domain, each with one Trp, were studied in the presence of the B1 domain of protein G, mutated to contain one Cys (K10C) and no Trp (W43Y). The concentrations of villin or SH3 were held constant at 30 μM. Concentrations of protein G were varied up to 0.84 mM. Higher concentrations could not be achieved in the experiment.

### Experiments

The Trp triplet state and its decay over time was then measured with two lasers in a pump-probe setup: a 10 ns pulse at 289 nm excites the triplet state while a continuous wave laser at 450 nm probes the population of the triplet state by transient absorption. We note that although Trp is much more sensitive to excitation than any other amino acid at these wavelengths, long-lived photophysics of tyrosine is also possible and the presence of four tyrosines in highly concentrated protein G leads to a significant background that must be subtracted, as shown in **Fig. S12**.

In Trp-Cys quenching, the Trp triplet decays exponentially with the quencher distance as

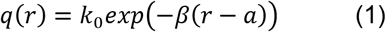

where *a* is the distance of closest approach determined here from simulation as a=3.5 Å (see below), *k*_0_=4.2×10^9^ s^−1^ is the decay rate at *a*, and *β* = 4 Å ^−1^ is the decay constant. The large value of β means that decay rates only remain significant up to distances of about a+5 Å. If interactions occur within this range, non-exponential decay curves are observed. Non-exponential behavior was first described for Trp and Cys amino acids immobilized in a sugar glass, which allowed determination of the quenching parameters above. Non-exponential kinetics is also observed when concentrations are high enough for molecules to diffuse in and out of close interactions on time scales shorter than the triplet state lifetime, *i.e*. <40 μs.

### Monte Carlo sampling of 1D potentials

We constructed Lennard-Jones-type potentials to create strong repulsion at close distances and weak attraction at the contact distance. Additional Gaussian functions were used to model kinetic barriers and kinetic traps at different distances (see **Fig 3**). The potential was mirrored at larger distances to create a finite-size system where concentration can be varied easily by adjusting the distance at which the potential is reflected. The potential was sampled via Monte Carlo sampling to create diffusive pseudo-dynamics without a meaningful time scale, but desired diffusion kinetics can be imposed by varying the Monte Carlo step size.

### Atomistic simulations

Molecular dynamics simulations in atomistic detail in the presence of explicit solvent were carried out for the experimentally studied systems. Because of computational limitations, the systems were simulated at about fivefold higher concentrations than in the experiments. Multiple simulations on μs time scales were carried out for each system and different force fields were tested (see summary in **Table S4**).

To compare the simulated-derived quenching decay curves directly with the experimental results, time-scale adjustments were necessary. First, the protein concentrations in the simulation were about fivefold larger than those in the experiment because the computational cost of long simulations at lower protein concentrations are too high and experiments at higher protein concentrations were not feasible. Faster diffusion in a less crowded environment may accelerate Trp-Cys quenching events, but fewer quencher proteins result in less frequent encounters. The experimental data collected at different protein G concentrations (**Fig. 2A**) indicates that the net effect is that at lower protein concentrations the quenching kinetics is slowed down. To extrapolate the dependence on concentration to the higher concentrations in the simulations, we carried out coarse-grained MD simulations at a range of concentrations between the concentrations in the experiments and simulations (**Fig. S13**). This allowed us to determine scaling factors for correcting the faster kinetics in the simulations due to the higher protein concentrations. Second, the TIP3P water model that was used in our simulations is known to underestimate viscosity, thereby accelerating diffusion, whereas the periodic boundaries applied in the simulations reduce the apparent diffusion over what would be obtained with infinitely large systems. Protein diffusion was calculated from mean square displacements (**Fig. S14**) with the resulting diffusion values before and after correction given in **Table S2**.

### Calculation of triplet state survival probability

From sampled probe-quencher distance time series, survival probability curves were calculated by evaluating **Eq. 2** based on the exponential quenching function in **Eq. 1** for the ‘distances’ sampled via Monte Carlo.

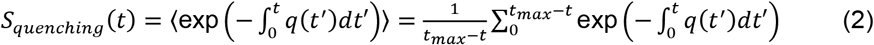

The triplet-to-singlet transition due to auto-bleaching was considered via **Eq. 3** with an auto-bleaching rate constant (k_0_) of 2.3 × 10^−5^ / ns (30).

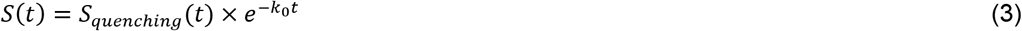

Additional details are given in Supplementary Material.

## Acknowledgments

Funding was provided by the National Science Foundation grants MCB 1817307 and by the National Institute of Health (NIGMS) grant R35 GM126948. Computer time was used on the Anton2 special-purpose supercomputer at the Pittsburgh Supercomputing Center (PSCA18053P).

