## Supplementary text, figures, and tables for "Characterizing Transient Protein-Protein Interactions by Trp-Cys Quenching and Computer Simulations"

**This PDF file includes:**

Supporting Text
- Sample preparation
- Trp-Cys quenching experiments
- Experimental data analysis
- Circular dichroism experiments
- Monte-Carlo sampling of 1D distance diffusion
- All-atom MD simulations
- Conformational state analysis
- Coarse-grained simulations
- Analysis of folding-unfolding equilibria

Figures S1 to S23
Tables S1 to S7
SI References

### Supporting Information Text

**Sample preparation.** Protein G (Lyophilized powder) and the peptides (villin WT, villin mutants: V10W, K33W, R15T+K30E R15T and SH3) were purchased from Biosynthesis in the form of white powder. No blocking groups were added at the termini.

Initial stock of protein G was prepared by dissolving in HEPES buffer (pH 7) and sonicating for 1 hr. 300 μM, 300 μL aliquots of the peptides were prepared by dissolving in HEPES buffer. Both protein G and peptide stocks were stored at -20°C.

Protein G was thawed and 2 ml of it was deoxygenated using $N_2O$ for 1 hr to scavenge solvated electrons created in the UV laser pulse and prevent $O_2$ quenching of the triplet. After deoxygenating an aliquot of the peptide was thawed and quickly added to the deoxygenated protein G in a cuvette and sealed where the final concentrations of the peptides are ~40μM.

**Trp-Cys quenching experiments.** Trp-Cys measurements were carried out on the prepared samples using pump probe spectroscopy at 20°C. Trp in peptides were excited to a long lived triplet state using a 10 ns laser pulse (pump beam) at 289 nm from a Nd:YAG laser and a 1-m Raman cell filled with 450 PSI of $D_2$ gas. The quenching of the Trp by the Cys in Protein G was observed as a decay by detecting the transient absorption in the triplet population using a 450 nm HeCd continuous laser (probe beam).

The probe beam was split to generate a reference beam and a sample beam that passed collinearly through the cuvette with the pump beam. The intensity of each beam was measured with a photodiode (New-Focus, Santa Clara, CA, Model 1621), combined in a differential amplifier (DA 1853A, LeCroy) with an additional stage of a 350-MHz preamplifier (SR445A, Stanford Research Systems), recorded with a digital oscilloscope (Tektronix TDS 3032) and stored on a computer. Measurements were obtained within about 15 min of addition of peptide to protein G. 6 measurements of 128 averages were obtained per sample. A baseline measurement of protein G without peptide was measured in the same way on the same day to reduce differences in instrument alignment.

**Experimental data analysis.** Each data set consists of two simultaneous measurements of 10,000 points 1 ns resolution and 1 μs resolution using two different oscilloscopes. The combined traces are then averaged in time to produce 128 points evenly spaced in logarithmically to best highlight decays on different time scales. For each peptide/protein G sample, the 6 measured decays were individually normalized at 100 ns to account for variations in laser intensity. The mean and standard deviation of the measurements was found point-by-point. The same procedure was used for protein G-only samples. The average trace for the protein G-only sample was subtracted from the peptide sample and the errors propagated by the sum of squares. Finally, the amplitude was normalized to be 0.88 between 100 and 150 ns and 0 at 360 μs to best match the atomistic simulation decays. These data and errors are plotted in Fig. 2. The derivative of the decays was found point-by-point by a linear regression of the normalized data over a window of 21 time points (in log($t$)). The slope and error of the fit are plotted in **Fig. 2**.

**Circular dichroism experiments.** Villin and SH3 samples were measured using a Jasco J-815 spectrometer. Samples were measured at room temperature using a 2 mm cuvette. Concentrations ranged from 12-26 μM and were normalized to match concentrations between samples. From the spectra, secondary structure percentages were estimated by the Jasco Multivariate Secondary Structure Estimation Program.

**Monte Carlo sampling of 1D distance diffusion.** The following 1D potential was constructed to model Trp-Cys distance diffusion:

$$V(r) = \varepsilon\left(\left(\frac{\sigma}{r}\right)^{12} - \left(\frac{\sigma}{r}\right)^{6}\right) + ae^{-(r-\mu)^2/w} + \frac{d\cdot k}{r}e^{-r/k} \quad \text{(S1)}$$

The potential combines a Lennard-Jones type contact potential with a Gaussian potential to model a kinetic barrier and a long-range Debye-Hückel type potential. The potential was sampled via Monte Carlo sampling at $kT$=1. Trial moves were taken randomly in positive or negative direction with a step size chosen randomly up until a maximum of 5 Å. For diffusion-matched runs, the maximum step size was adjusted as a function of *r* to model different systems (see **Table S5**). Typical sampling runs involved $10^7$ trial moves. A finite system size was modeled by wrapping distance values around when a given maximum size was exceeded, effectively mirroring the potential at the maximum size. Potential parameters were adjusted to match probability distributions extracted from the atomistic simulations and then further modified to explore the effect of potential features on the resulting quenching curves (see **Table S5**). When the long-range potential was used (*i.e.* $d \neq 0$), a value of $k$ = 10 Å was used as the screening term.

A Jupyter notebook implementation of the Monte Carlo sampling program is available on github at https://github.com/feiglab/mc-trpcys

**All-atom MD simulations.** Protein structures were retrieved from the Protein Data Bank (PDB) with PDB IDs of 1VII, 2A37, and 1GB1 for villin WT, SH3 T22G, and protein G structures, respectively. Missing atoms were added using the MMTSB Tool Set (1). Structures for villin mutants (V10W, K33W, and R15T+K30E) were also prepared with the MMTSB Tool Set. The list of probes and quencher (Protein G) proteins are summarized in **Table S3**. One probe and nine quencher protein structures were randomly oriented and placed in a cubic box with a width of 144.048 Å, corresponding to concentrations of 0.56 and 5.0 mM for the probe and quencher proteins, respectively. The rest of the cubic box was filled with explicit water molecules. Finally, some water molecules were randomly selected and replaced with either sodium or chloride ions to achieve overall neutral systems with an excess ion concentration of 25 mM. The constructed all-atom simulation systems were composed of around 297,000 atoms.

Proteins were described with CHARMM force fields. Four CHARMM force fields variants were considered: (1) CHARMM 36 (c36) (2), (2) CHARMM 36m (c36m) (3), (3) CHARMM 36 with scaling of water interactions (c36+water) (4), and (4) CHARMM 36mw (c36mw) (3). The water molecules were described with the CHARMM version of TIP3P (5).

MD simulations were carried out using Anton2 (6) and OpenMM (7). For Anton2 simulations, simulation systems were initially equilibrated using NAMD (8). The systems were locally minimized for up to 1,000 steps first. The minimized systems were then equilibrated via a series of Langevin dynamics simulations with a friction coefficient of 1.0/ps. A 2-fs integration time step was used, while hydrogen-containing bonds were kept rigid using the SHAKE algorithm. Equilibration consisted of gradual heating from 20 to 298.15 K with an increment of 20 K for every 10 ps. During the simulations, the size of the simulation boxes remained unchanged as the NVT ensemble was applied, and every Cα atoms were restrained with positional harmonic restraints with a force constant of 1.0 kcal/mol/Å$^2$. Then, the systems were further equilibrated in the NPT ensemble at 1 bar with a Nose-Hoover Langevin barostat. Another round of Langevin dynamics simulations were carried out for 1 ns with the same restraints, and the applied restraints were gradually loosened with followed 2.5 ns-long simulations with force constants of 0.5, 0.1, 0.05, 0.01, and 0.0 kcal/mol/Å$^2$. Files for the equilibrated systems were converted into Anton2 input files, and simulations for production were performed on Anton2. The simulations were carried out using a multi-time step integrator (RESPA) with a 2.5 fs integration time step. Bonded and near-range non-bonded interactions were evaluated for every step while far-range interactions were evaluated for every three steps. Temperatures were maintained at 298.15 K using a Nose-Hoover thermostat with an interval of 60 fs, and pressure was set to 1 bar controlled by a Martyna-Tobias-Klein (MTK) barostat with an interval of 1.2 ps. Simulations were continued for at least 3.0 μs, and snapshots of the simulations were recorded for every 240 ps. Simulation details for each system are given in **Table S4**.

Simulations using OpenMM were carried out using a similar procedure with minor differences: Simulation systems were minimized using the l-BFGS-b minimizer for up to 500 steps. The minimized systems were subjected to similar equilibration steps. Langevin dynamics simulations were facilitated with a friction coefficient of 0.01/ps and a 2-fs integration time step with the SHAKE algorithm. The same restraints were applied to every Cα atoms with a force constant of 0.5 kcal/mol/Å$^2$. Once the systems were heated up through the same heating schedule, they were equilibrated for 1 ns in the NPT ensemble. Pressure was controlled to 1 bar using a Monte Carlo barostat. Then, the applied restraints were gradually loosened using the same schedule used for the equilibration step for Anton2 simulations except for an additional step with a force constant of 1.0 kcal/mol/Å$^2$. In the production stage, Langevin dynamics simulations were performed with a 0.01/ps friction coefficient and a 2.5 fs integration time step and continued for at least 2.4 μs. Snapshots were stored at every 240 ps.

The minimum heavy-atom distances were evaluated between the indole ring of the Trp sidechain in a probe and the sulfur atom of Cys in the quenchers for every snapshot at different time points *t* to obtain *r* in **Eq. 1**. Triplet survival probabilities were then calculated as for the Monte Carlo sampling data based on **Eq. 2**.

The nominal simulation time scale was corrected to account for the lower viscosity of the TIP3P water molecules and periodic boundary condition (PBC) artifacts by applying the factor given in **Eq. S2**.

$$f_{TIP3P,PBC} = \frac{\eta_{w,exp}}{\eta_{TIP3P}} \times \frac{D_{t,MD}}{D_t} \tag{S2}$$

The lower viscosity of TIP3P was considered via the ratio between the experimental viscosity of pure water (0.89 cP) over TIP3P water (0.35 cP). The PBC artifact was considered via the ratio between the uncorrected and corrected translational diffusion coefficients for a given system. Translational diffusion coefficients $D_{t,MD}$ were obtained from mean square displacements (MSD) of the centers of mass of the proteins (**Fig. S14**) according to the Einstein relation (**Eq. S3**) based on linear fits for lag times from 1 to 100 ns.

$$D_{t,MD} = \frac{MSD(\tau)}{6\tau} \tag{S3}$$

A correction term for system-size effects due to PBC was then added according to **Eqs. S4** and **S5** (9). The hydrodynamic radii ($R_h$) of the proteins were obtained using HYDROPRO, and the average periodic box width was used for the system size (*L*).

$$D_t = D_{t,MD} + D_{t,PBC} \tag{S4}$$

$$D_{t,PBC} = \frac{kT}{6\pi\eta L}\left(\xi - \frac{4\pi R_h^2}{3L^2}\right) \tag{S5}$$

The shear viscosity ($\eta$) of our crowded system was estimated by **Eq. S6** according to the crowder volume fraction.

$$\eta = \eta_{TIP3P}\left(1 + 2.5\,\frac{\sum_k \frac{4}{3}\pi R_{h,k}^3}{L^3}\right) \tag{S6}$$

Another time correction factor was introduced to account for different protein concentrations in the simulations and experiments. To estimate, how diffusion is affected by concentration we used information from coarse-grained (CG) simulations. The detailed procedure of the CG simulations is described below. We assumed that the concentration of proteins would not alter the shape of survival probability curves significantly, but only shift them in time scale. Survival probability curves at various protein concentrations were evaluated using **Eq. 1** and **2** but with kinetic constants for the CG model. The kinetic constant parametrization for the CG model is described below. We note that the time correction factor (**Eq. S2**) was not applied because the simulations effectively used implicit solvent and because we are extracting only relative rates of diffusion at different concentrations from the CG model. The quenching curves from the CG simulations were rescaled to match the experimental data at concentrations of 0.5 mM and 0.8 mM where experimental data is available (**Fig. 2**). This allowed use to estimate the slow-down in kinetics at the higher concentrations in the all-atom simulations. Consequently, we obtained the following time scale correction factors for different system to allow for direct comparison between the simulations at higher concentrations vs. the experimental data: 11.096 for villin WT, 9.019 for villin V10W, 10.796 for villin K33W, 8.350 for villin R15T+K30E, and 11.447 for SH3.

**Conformational state analysis.** The conformational sampling of the probe proteins (villin and SH3) were explored via MD simulations, followed by Markov state modeling (MSM) analysis (10). For each probe protein, starting from its native state conformation, ten replicas of 200 ns-long MD simulations were carried out. The sampled conformations were then clustered to define conformational states. In a second round of simulations, starting from newly found conformations, another set of MD simulations were carried out. MD simulations and clustering until the MSM model reached convergence, typically after three or four iterations with 40–50 μs of simulation time in total.

For each initial conformation, MD simulations were performed in an explicit water box. One probe protein structure was located at the center of a simulation box, and the CHARMM version of TIP3P water molecules filled the box with at least 9 Å from any protein atom. The system was described with the CHARMM 36m force field. The charge of the system was neutralized by either sodium or chloride ions. The prepared system was locally minimized for 500 steps using the l-BFGS-b algorithm and equilibrated using Langevin dynamics simulations for 1 ns. It was gradually heated up to 298.15 K in the NVT ensemble and was further equilibrated at 1 bar in the NPT ensemble with a Monte Carlo barostat. During the equilibration steps, all hydrogen-containing bonds were kept rigid using the SHAKE algorithm, and protein structures were restrained by applying harmonic restraints on every Cα atom with a force constant of 0.5 kcal/mol/Å$^2$. For an equilibrated system, ten replicas of 200 ns-long Langevin dynamics simulations were performed with a friction coefficient of 0.01/ps. Snapshots are recorded at every 100 ps.

Markov state models were built from the simulation trajectories using PyEMMA (11). The distance matrices for Cα atoms were extracted from the trajectories and used to distinguish conformations. The dimension of the matrices was reduced to two using tICA (time-structure independent component analysis) with a lag time of 48 ns, which represented good Markovian behavior. Then, conformations were clustered into 200 clusters to define microstates using the k-means clustering algorithm with Euclidean distance in the reduced dimension. The microstates were lumped into nine macrostates based on a lag time of 48 ns. To ensure that sampling for all villin mutants can be mapped onto the same reaction coordinates and state definitions, trajectories for all mutants were used together to determine tICA coordinates. Then, free energy landscapes were evaluated separately for each villin mutant and mapped onto the common low-dimensional tICA space. In addition, villin conformations from simulations in the presence of quencher protein were also mapped onto the same reaction coordinates to compare the conformational sampling between the two sets of simulations.

**Coarse-grained simulations.** Residue-level coarse-grained MD simulations were carried out to explore quenching kinetics at lower concentrations and for a wider range of conformational states for the probe protein than what was feasible via all-atom modeling. In the coarse-grained model, a protein residue was represented as a spherical particle. The potential energy function is given in Eq. S7.

$$U_{total} = \frac{1}{2}k_{bond}(l-l_0)^2 + \frac{1}{2}k_{angle}(\theta-\theta_0)^2 \\ +4\left(\epsilon_{i,j}+\epsilon_{cation-\pi,i,j}\right)\left[\left(\frac{\sigma_{i,j}}{r_{i,j}}\right)^{10}-\left(\frac{\sigma_{i,j}}{r_{i,j}}\right)^{5}\right]+\frac{A_iA_j+(A0_i+A0_j)}{r_{i,j}}e^{-\frac{r_{i,j}}{\kappa}} \quad \text{(S7)}$$

Bonds between two neighboring residues ($l$) were described by a harmonic bond potential with a spring constant ($k_{bond}$) of 41.84 kJ/mol/Å$^2$ and an equilibrium bond length ($l_0$) of 3.8 Å. Bond angles for three consecutive residues ($\theta$) were described by a harmonic bond angle potential with a spring constant ($k_{angle}$) of 4.184 kJ/mol/rad$^2$ and an equilibrium bond angle ($\theta_0$) of 180°. Non-bonded interactions were evaluated for every residue pair with a distance ($r_{i,j}$) except for bonded residue pairs with a distance cutoff of 30 Å. For the short-range 10-5 Lennard-Jones potential, the radius of each amino acid particle ($\sigma_i$) was determined from the radius of a sphere of equivalent volume of the corresponding amino acid, and the sum of radii of two interacting particles ($\sigma_{i,j}$) was used for the calculation. The strength of short-range attraction ($\varepsilon_i$) was set to 0.40 and 0.41 kJ/mol for polar (Arg, Asn, Asp, Cys, Gln, Glu, His, Lys, Ser, and Thr) and non-polar (Ala, Gly, Ile, Leu, Met, Phe, Pro, Trp, Tyr, and Val) residues, respectively. In addition, cation-$\pi$ interactions between Arg/Lys and Phe/Tyr/Trp were augmented by a parameter, $\varepsilon_{\text{cation-}\pi}$, of 0.30 kJ/mol. For the long-range

Debye-Hückel potential, $A_i$ and $A0_i$ values were introduced for every residue in the unit of kJ/mol. $A_i$ was calculated according to **Eq. S8** for every residue according to its net charge ($q_i$). (12)

$$A_i = sign(q_i)\sqrt{0.75|q_i|} \quad \text{(S8)}$$

$A0_i$ was set to 0.05 for polar and 0 for non-polar residues to account for repulsive interactions between polar residues due to solvation. The ion concentration was implicitly described by the Debye-Hückel screening length ($\kappa$) We used the same ion concentration that we used for the all-atom simulations, 25 mM, where the corresponding screening length is around 12 Å.

While the CG model was initially designed for intrinsically disordered peptides, we applied additional distance restraints to maintain folded structures. A force constant of 2.092 kJ/mol/Å$^2$ to Cα atom pairs that were separated by two or more residues and that had distances below 20 Å.

The CG model was implemented in OpenMM (7) to take advantage of GPU acceleration and simulated via Langevin dynamics. It was validated for the modeling of our systems by comparing radial distribution functions from the CG simulations with those using all-atom MD simulation trajectories (**Fig. S15**). Even though heights of peaks for probe-quencher interactions were lower than those in all-atom simulations, the overall interaction preference trends across various probe proteins were consistent with the all-atom simulation data. There was also good agreement in the peak locations. Furthermore, the CG simulations reproduced the radial distribution functions for quencher-quencher interactions well. Therefore, we concluded that the coarse-grained model was valid for describing our systems.

Kinetic constants of the Trp-Cys quenching for the CG model were determined by matching the derivative of the survival probability curve for the coarse-grained model with that for the all-atom model for the villin WT system. A grid search was performed with $\beta$ values of 1, 2, 4, and 6 / Å, $q_0$ values of $10^{-3}$, $10^{-2}$, $10^{-1}$, 1.0, and 4.2 / ns. For $a_0$, we tested the sum of radii for Trp and Cys CG particles (~6.04 Å) with offsets range from -2 to +2 Å with a space of 0.5 Å. From the search, we determined that a parameter set with a $\beta$ of 2 / Å, a $k_0$ of 4.2 / ns, and a distance offset of -2.0 Å ($a_0$=4.04 Å) resulted in the best agreement for the derivative curve with respect to the reference all-atom simulation.

To investigate the protein concentration dependence, coarse-grained MD simulations were carried out at different quencher protein concentrations, and survival probability curves were calculated and compared with the experimental results as described above. The simulation systems were prepared with a probe protein concentration of 0.05 mM and a range of quencher protein concentrations (0.5, 0.8, 2.0, and 5.0 mM). This was realized by one probe together with 10, 16, 40, or 100 quencher proteins in a cubic box with a width of 321.435 Å, respectively.

Another set of coarse-grained MD simulations was performed to evaluate the survival probability decay as a function of conformational state of a probe protein. For each state of a probe protein, a representative structure from the MSM analysis was utilized as an initial conformation for the coarse-grained simulations. For an initial conformation, a coarse-grained simulation system was prepared in the same condition (*e.g.,* simulation box size and the number of proteins) that was used for the all-atom simulations. For both sets of simulations, three replicas of Langevin dynamics simulations were carried out at 298.15 K for 60 µs with an integration time step of 20 fs, and snapshots were saved every 100 ps.

Analogous to the calculation of the survival probability for the all-atom model, a time correction factor was introduced to account for faster diffusion in the coarse-grained simulation:

$$f_{CG}(c) = \frac{D_{t,CG}(c)}{D_{t,est}(c)} \quad \text{(S9)}$$

We measured from the all-atom simulation that bigger protein clusters tend to diffuse slower. (**Fig. S16**) The tendency of transient cluster formation varied at different protein concentrations, the translational diffusion coefficients at a concentration, $D_{t,est}(c)$ can be estimated by a weighted sum of coefficients for each cluster size with a weight of the corresponding cluster size population:

$$D_{t,est}(c) = \sum_n p'_{CG}(n;c) \times D_t(n) \quad \text{(S10)}$$

However, transient cluster formation in the coarse-grained simulation was not as big as that in the all-atom simulations (**Fig. S17**). Thus, the population of each cluster size should be adjusted. We assumed that the tendency of changes in cluster size populations was the same as in the one probe and nine quencher protein system across different concentrations:

$$p'_{CG}\,(n;c) = p_{CG}\,(n;c) \times \frac{p_{AA}\,(n;c^{o})}{p_{CG}\,(n;c^{o})} \qquad \text{(S11)}$$

The time correction factor was multiplied to the simulation time to adjust for the faster diffusion in the coarse-grained simulations.

**Analysis of folding-unfolding equilibria.** To check that proteins remain folded, we carried out CD spectroscopy for villin, SH3, and protein G (**Fig. S18**). The measured CD spectra for villin indicates largely α-helical secondary structures with some differences for different mutants that indicate mostly folded structures but with some variations in secondary structure content (**Table S6**) and with similar helical secondary structure contents as estimated from the simulations (**Table S7**) where villin remained largely folded. Therefore, we conclude that the mutations only induce minor structural perturbations and that all villin variants are largely present as globular three helix bundles as the wild type.

The SH3 spectra resemble published spectra for drkN SH3 domains and indicate a mixture of folded beta and disordered conformations (13). We used here the T22G mutant, that is expected to be more stable than the wild-type sequence, but CD spectra indicate less stability intermediate between the wild-type and previously recorded CD spectra for the T22G mutant. Quantitative analysis suggests that some of the SH3 structures may have partially lost their native structure, since the sheet content estimated from the CD spectrum is only half (23%, **Table S6**) than what is found in the simulations of the natively folded structure (42%; **Table S7**).

The presence of partially unfolded states may affect the interpretation of the Trp-Cys quenching results. To address this point, we carried out additional simulations to sample extended ensembles of villin and protein G, including partially non-native states (see Supplementary Material). A number of different states were identified (**Figs. S19-S23**) based on Markov State Modeling analysis. The different states were then subjected to coarse-grained simulations to estimate the effect on altered quenching curves (**Figs. S19-S23**). Essentially, we find little difference in the quenching curves with only a few states showing noticeable differences. Therefore, we conclude that even in the presence of some partial deterioration of the native structures, the Trp-Cys quenching curves are not affected significantly and that the general observations remain valid.

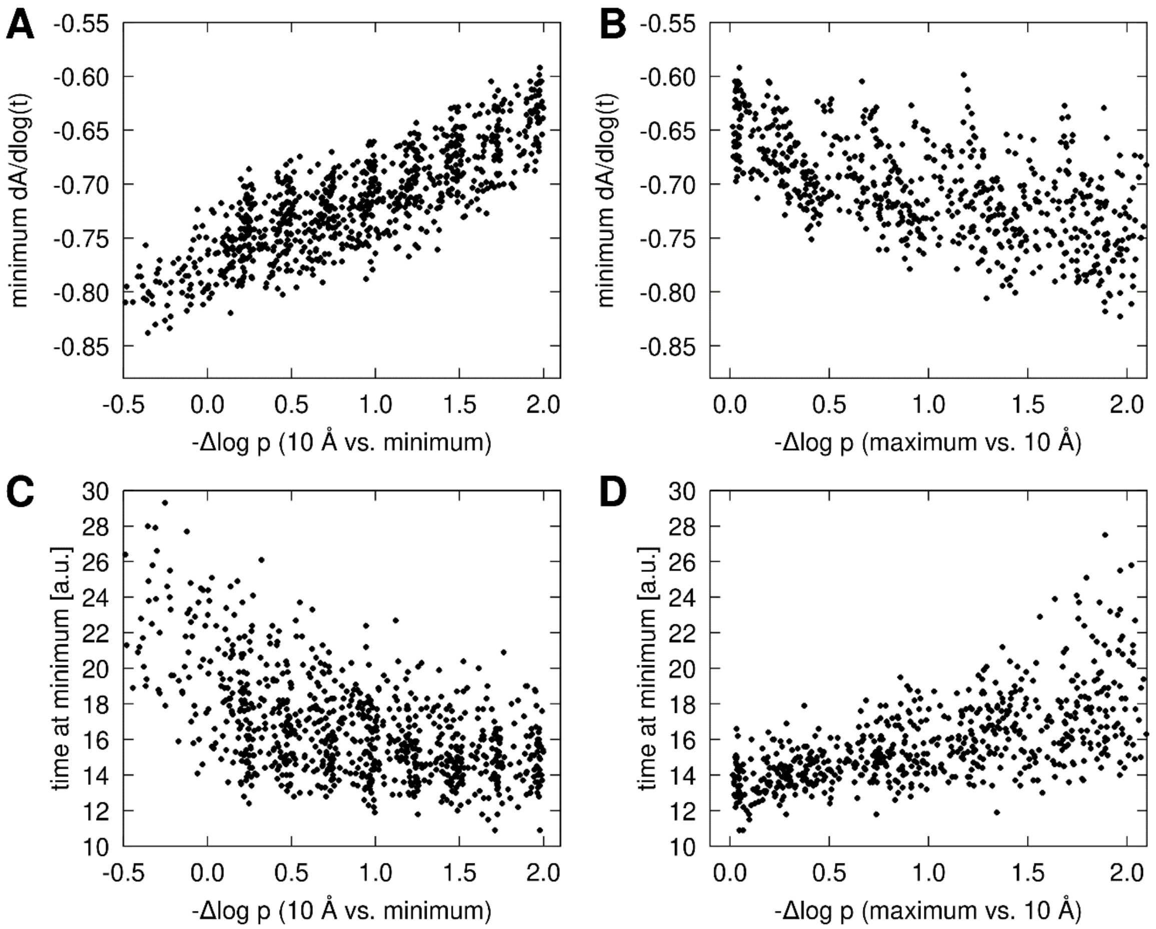


**Figure S1**. Dependence of quenching curve derivative minimum on 1D potential features for the minimum value (A and B) and the time at which the minimum is reached (C and D) as a function of the difference in -log p between 10 Å and the contact minimum and between -log p at the maximum barrier height and the value at 10 Å. The potential was varied in terms of $\varepsilon$ (1-8), the barrier height $a$ (0-3), the barrier location $\mu$ (4.5-6.0 Å) and the barrier width $w$ (0.2-1.2 Å). $\sigma$ was fixed at 3.28 Å (see **Eq. S1**). No long-range attraction or repulsion was considered.

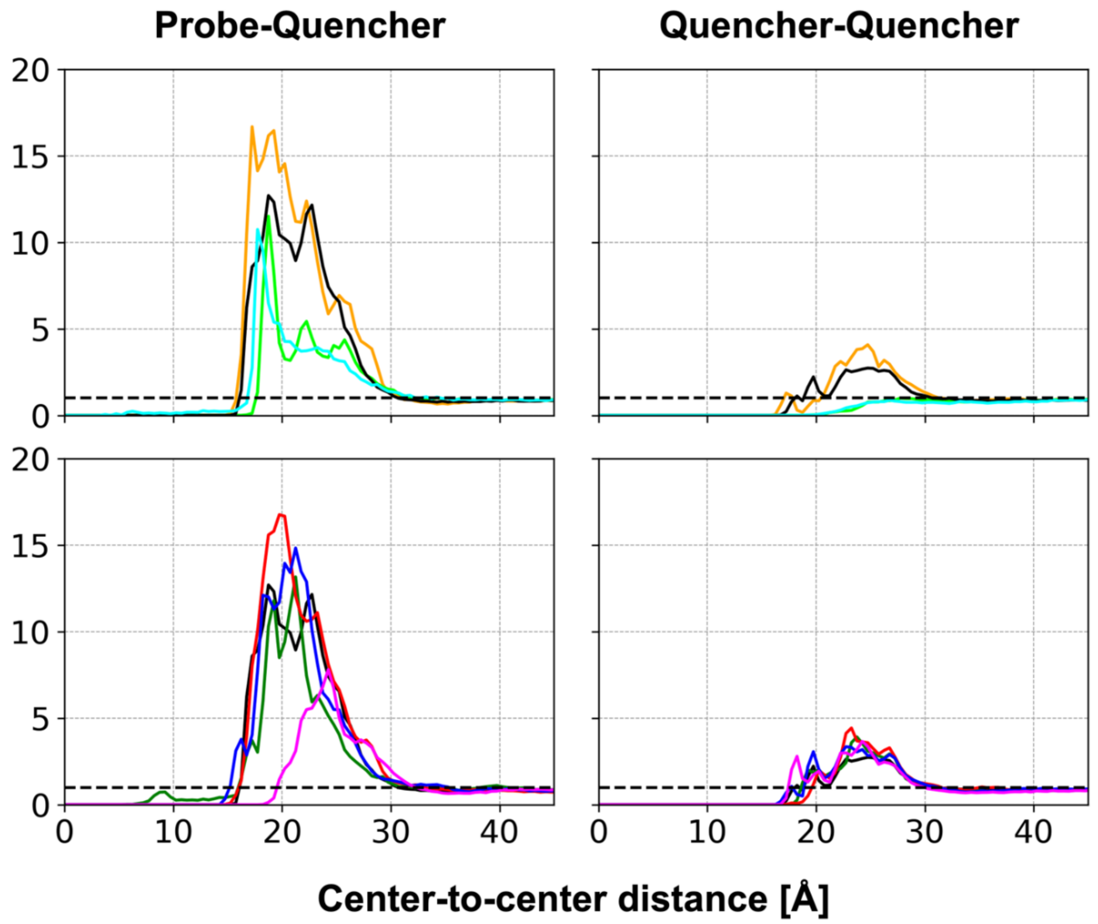


**Figure S2**. Radial distribution functions between proteins. (Top) comparisons between force fields for the villin WT system: c36 (orange), c36m (black), c36+water (lime), and c36mw (cyan). (Bottom) comparisons between different probe proteins: villin WT (black), villin V10W (red), villin K33W (blue), villin R15T+K30E (green), and SH3 (magenta). Black dashed lines are overlaid for g(r)=1.

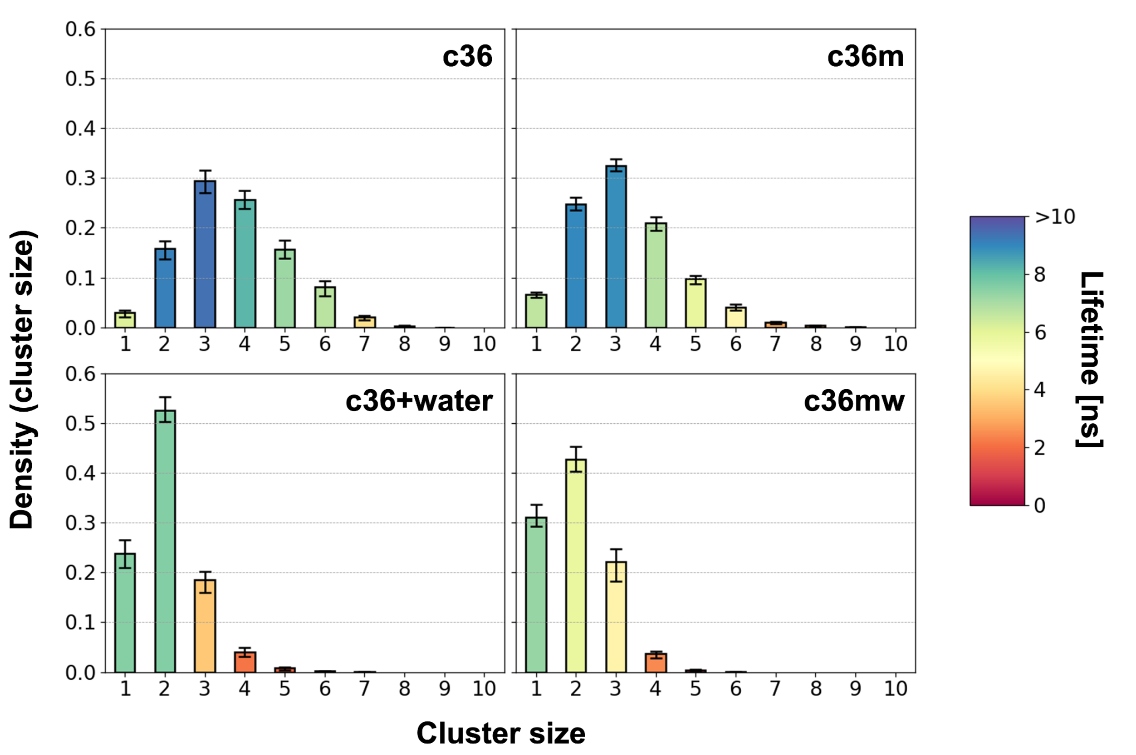


**Figure S3**. Cluster size distributions and corresponding lifetimes for villin WT with different force fields. The cluster size distributions are shown as bar charts with error bars for standard errors. Each bar is colored according to its cluster size lifetime, and the lifetime refers to the mean first passage time (MFPT) to other cluster sizes.

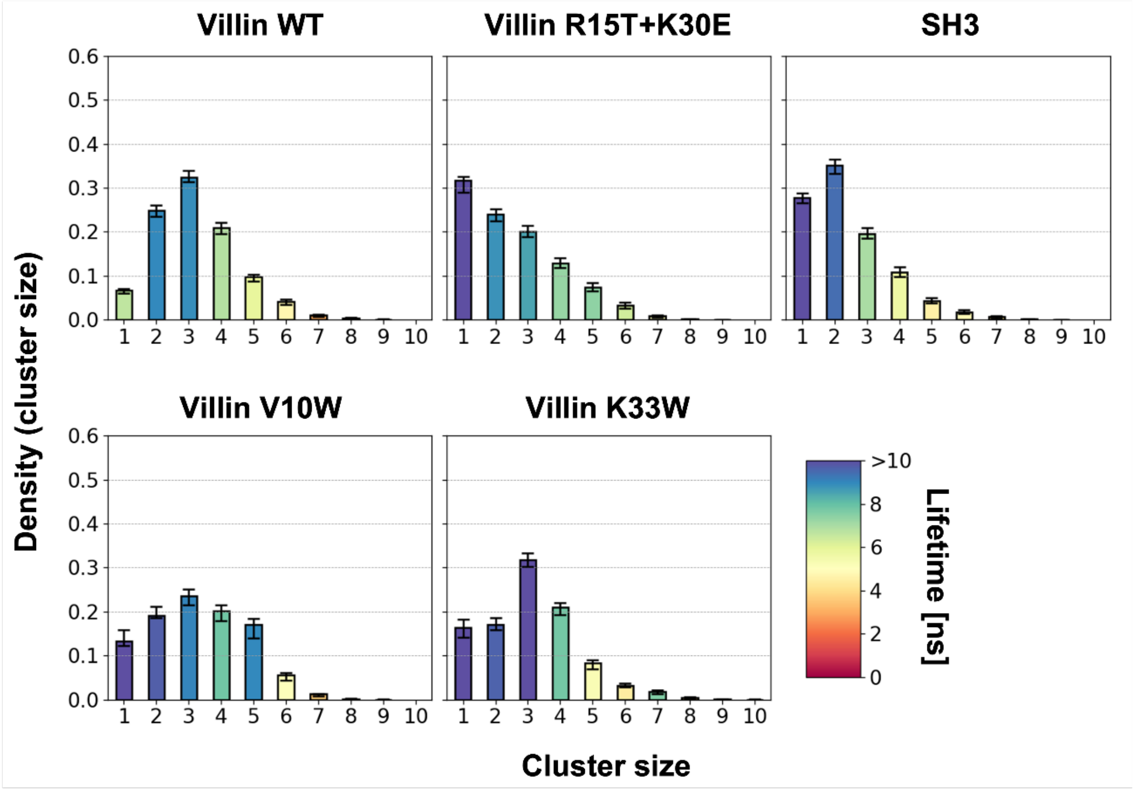


**Figure S4.** Cluster size distributions and corresponding lifetimes for probe protein in different systems. The cluster size distributions are shown as bar charts with error bars for standard errors. Each bar is colored according to its cluster size lifetime, and the lifetime refers to the mean first passage time (MFPT) to other cluster sizes.

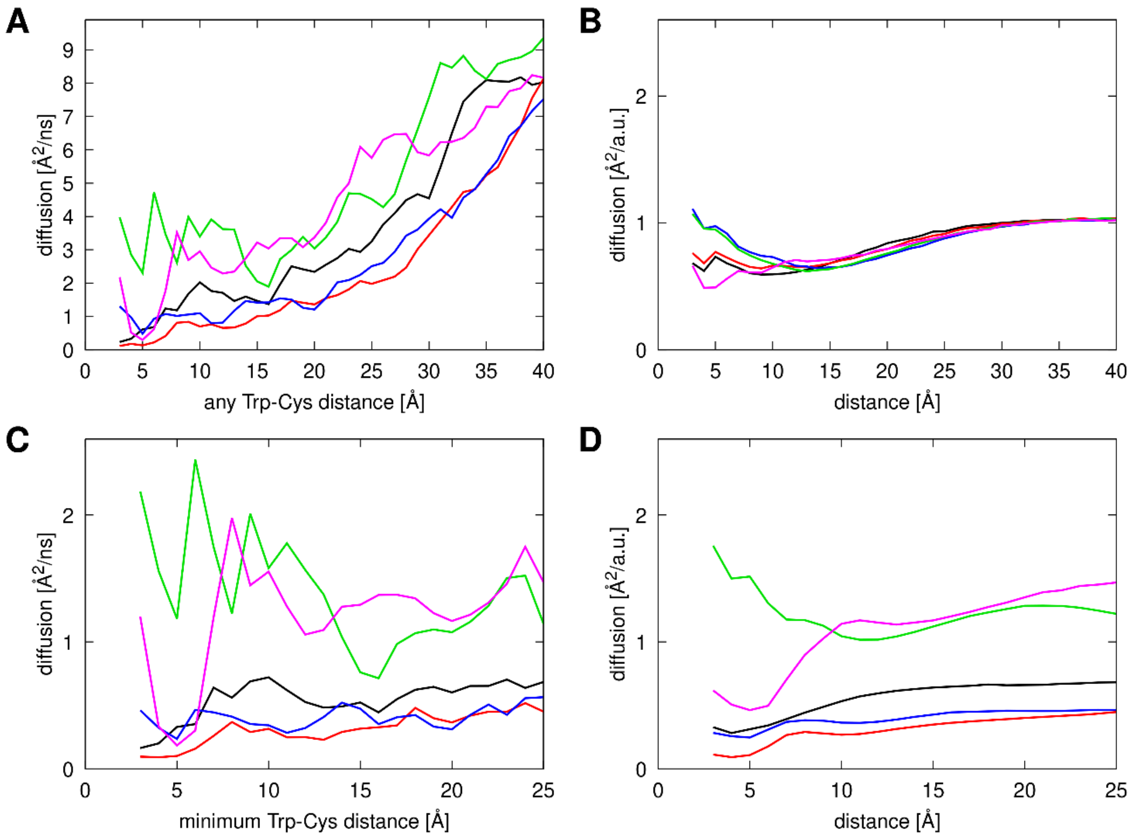


**Figure S5**. Diffusion in Trp-Cys distance variable from atomistic simulations (A,C) and Monte Carlo-based 1D potential sampling (B,D) for villin variants and SH3 (colored as in Fig. 2). Diffusion based on individual protein G-villin distance fluctuations is shown in A, diffusion based on minimum distance of any protein G to villin is shown in C. Monte-Carlo based sampling without adjustment is shown in B, and after distance-based, system-dependent scaling of the maximum Monte Carlo step to match the atomistic data is shown in D (see Methods). Diffusion was calculated based on linear fits to MSD curves up to 20 ns. Distance-dependent diffusion was obtained by averaging separate MSD curves depending on the initial Trp-Cys value for a given time interval for which MSD was calculated.

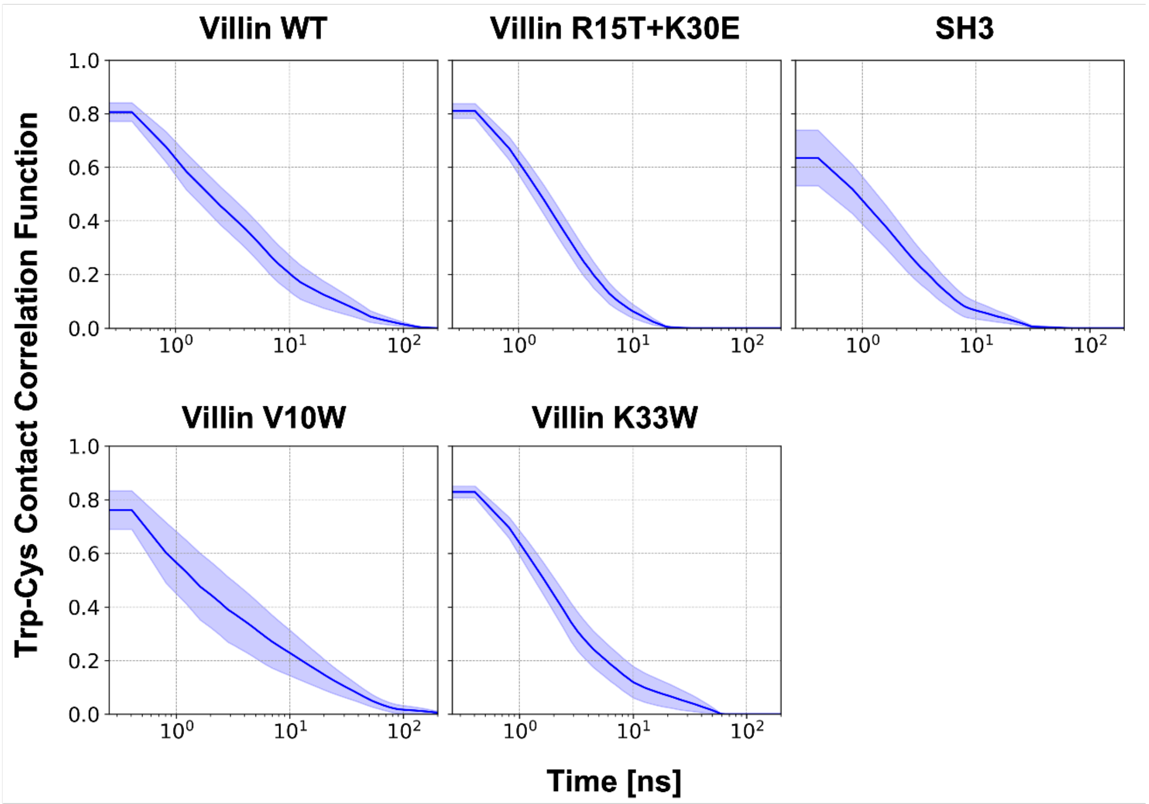


**Figure S6.** Trp-Cys contact correlation function. Simulation trajectories with CHARMM 36m force field were used for the analysis. The contact was defined If the minimum heavy-atom distance between a tryptophan and a cysteine was closer than 5 Å. The lifetime for Trp-Cys contact from fitting of the contact correlation function was 20.7, 3.9, 26.5, 10.4, 5.6 ns for villin WT, R15T+K30E, V10W, K33W, and SH3, respectively.

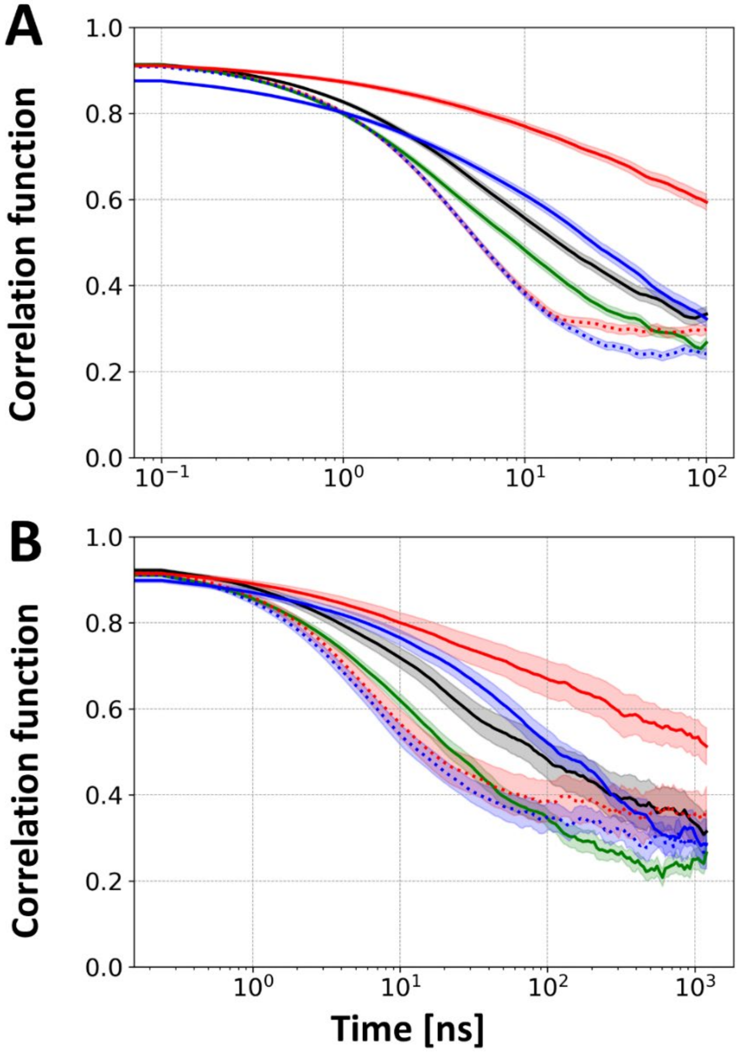


**Figure S7**. Time correlation function of Trp (Tyr) vector. The vector was defined as a unit vector of the residue from the Cα atom to the center of mass of the sidechain except for the Cβ atom after structure alignment with respect to the initial conformation using backbone and Cβ atoms of the residue and adjacent residues. Time correlation function was evaluated by calculating the ensemble average of the P2 Legendre polynomial of the inner product between the vectors. It was evaluated using all-atom simulation trajectories of without (A) and with (B) protein G systems: villin WT (black), V10W (red), K33W (blue), and R15T+K30E (green). Data for the Trp vector and the Tyr24 vectors for single mutants are shown as solid and dotted lines, respectively.

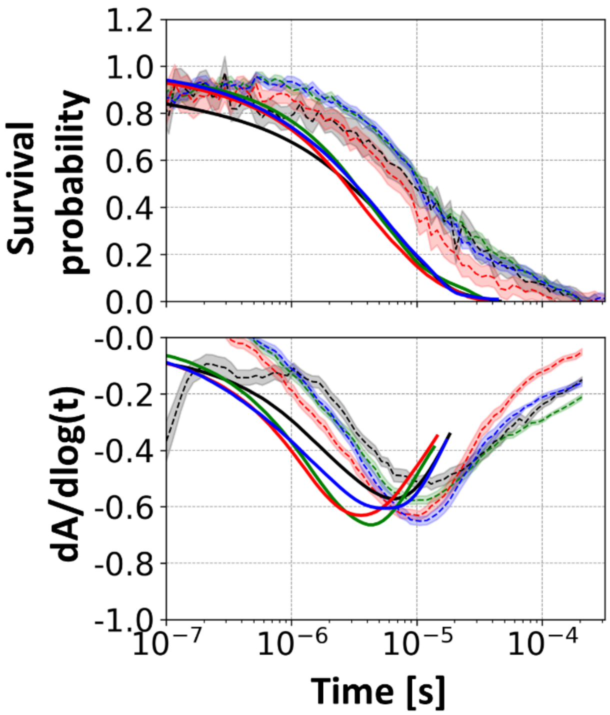


**Figure S8**. Comparison of survival probability decays between experimental measurements and simulated data as in Fig. 4, but with V10W and K33W quenching curves calculated with respect to hypothetical quenching at the 24 position instead of the actual tryptophan at residues 10 and 33, respectively.

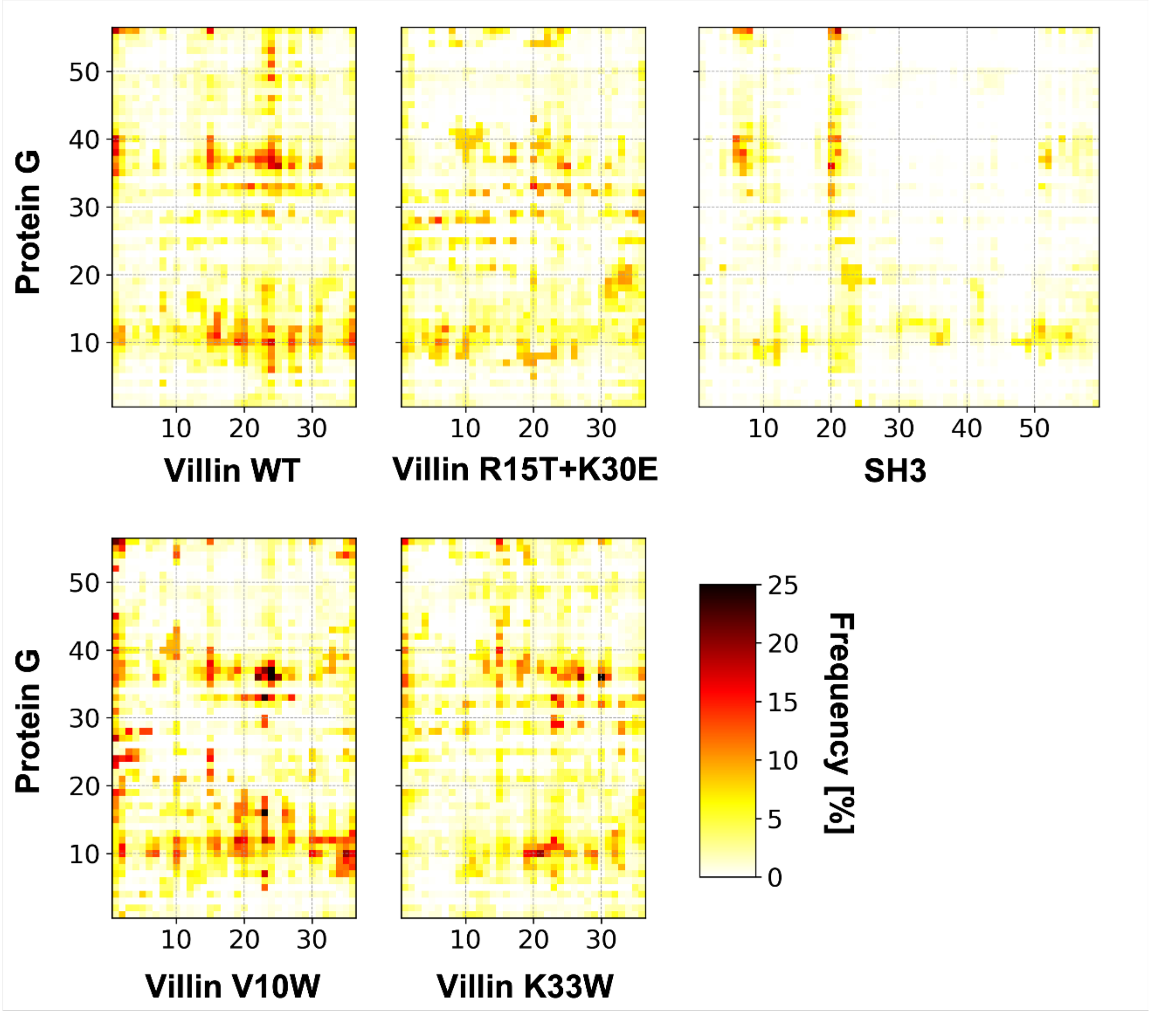


**Figure S9.** Inter-residue contact frequencies between probe and quencher proteins. Inter-residue contacts were defined as residue pairs whose inter-atomic distances between them were closer than 5 Å. More frequent contacts are depicted in dark reddish colors.

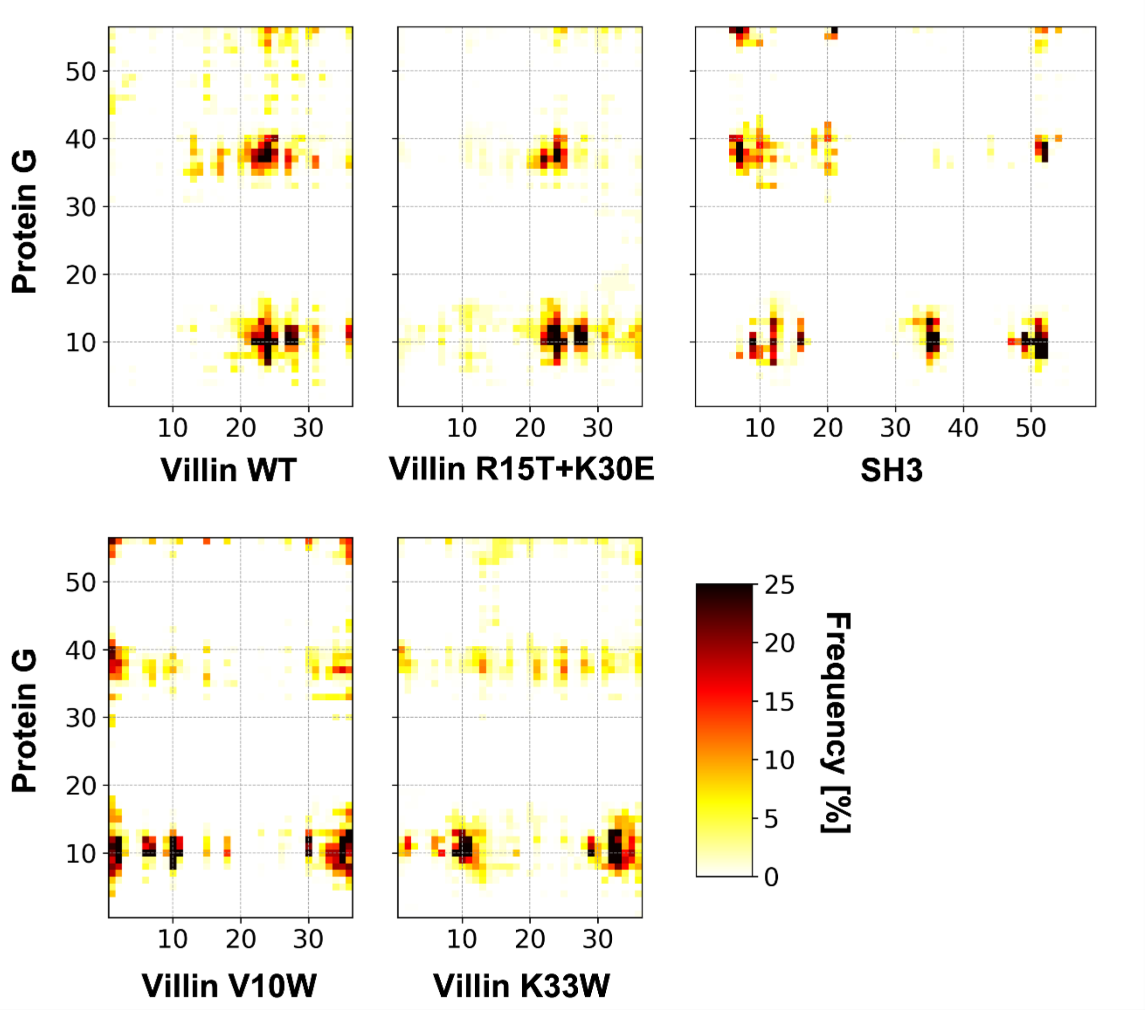


**Figure S10.** Inter-residue contact frequencies between probe and quencher proteins as in **Fig. S9** but only at the time of Trp-Cys quenching contact.

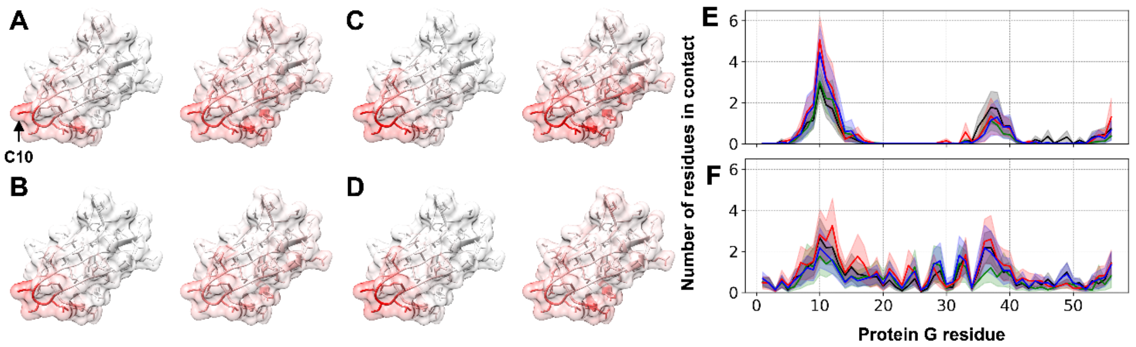

**Figure S11**. Residue-wise contacts between protein G and villin variants, projected onto the protein G (A-D) and as a function of protein G residue index (E, F). Surface projections are shown at the time of Trp-Cys quenching contact (left) and at any time of contact (right) for wild-type villin (A), the R15T+K30E mutant (B), the V10W mutant (C), and the K33W mutant (D). The location of the Cys residue is indicated by arrows. Contacts per frame vs. residue index are shown at the time of quenching contact (E) and at any time of contact (F) with different variants colored as in **Fig. 2**. Shaded areas indicate standard errors. Contacts were defined by residue pairs whose inter-atomic distances were closer than 5 Å.

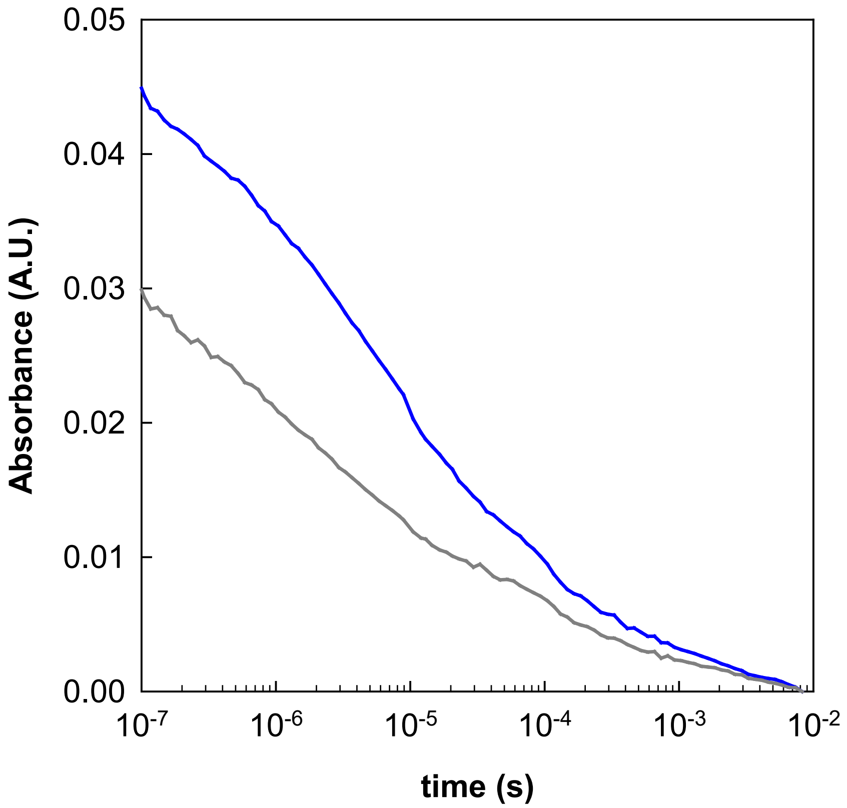


**Figure S12**. Example of protein G background subtraction in Trp-Cys measurements. Samples of protein G (840 μM) with (blue) and without (grey) 80 μM K33W villin were prepared and measured on the same day. Each trace is average of six measurements. The protein G only sample is subtracted from the K33W/protein G without adjustment, then normalized as shown in **Fig. 2**.

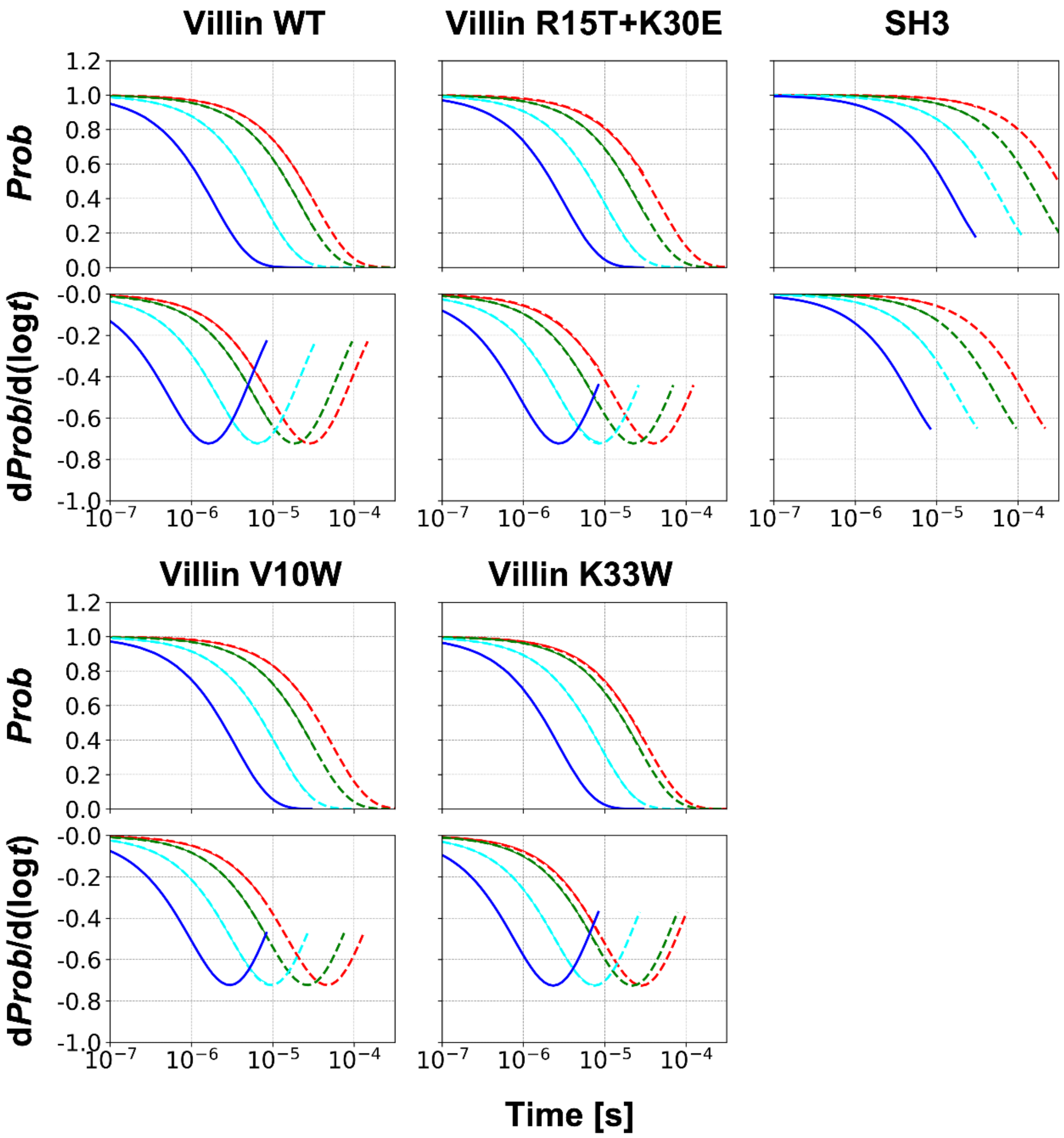


**Figure S13.** Survival probability as a function of quencher concentration from coarse-grained MD simulations. Simulations at concentrations of 0.5, 0.8, 2.0, and 5.0 mM are shown as red, green, cyan, and blue lines, respectively. Rescaled quenching curves for each concentration by applying time rescaling factors to the curve for 5.0 mM are shown as dashed lines in the corresponding colors.

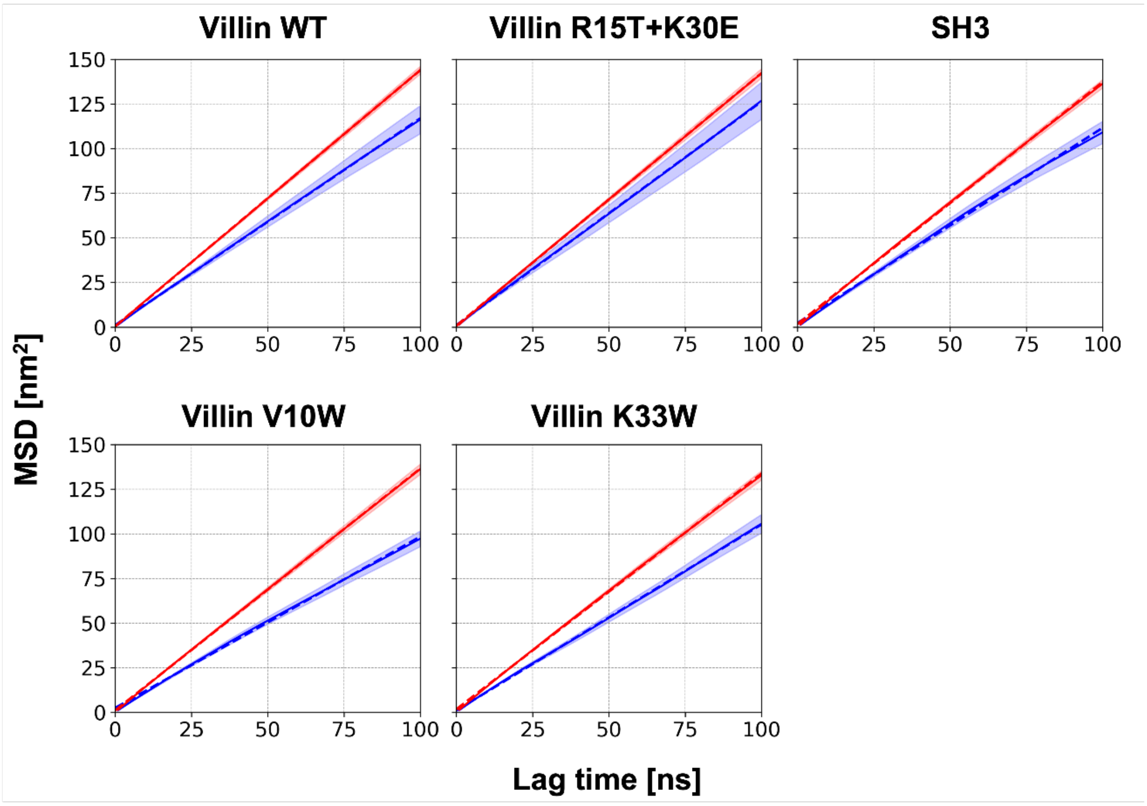


**Figure S14**. Mean square displacement (MSD) versus lag time for probe (blue) and quencher (red) proteins. Average and standard error of MSD are shown with a solid line and transparent shade. The linear-fit used for determining translational diffusion coefficients is shown as dashed lines.

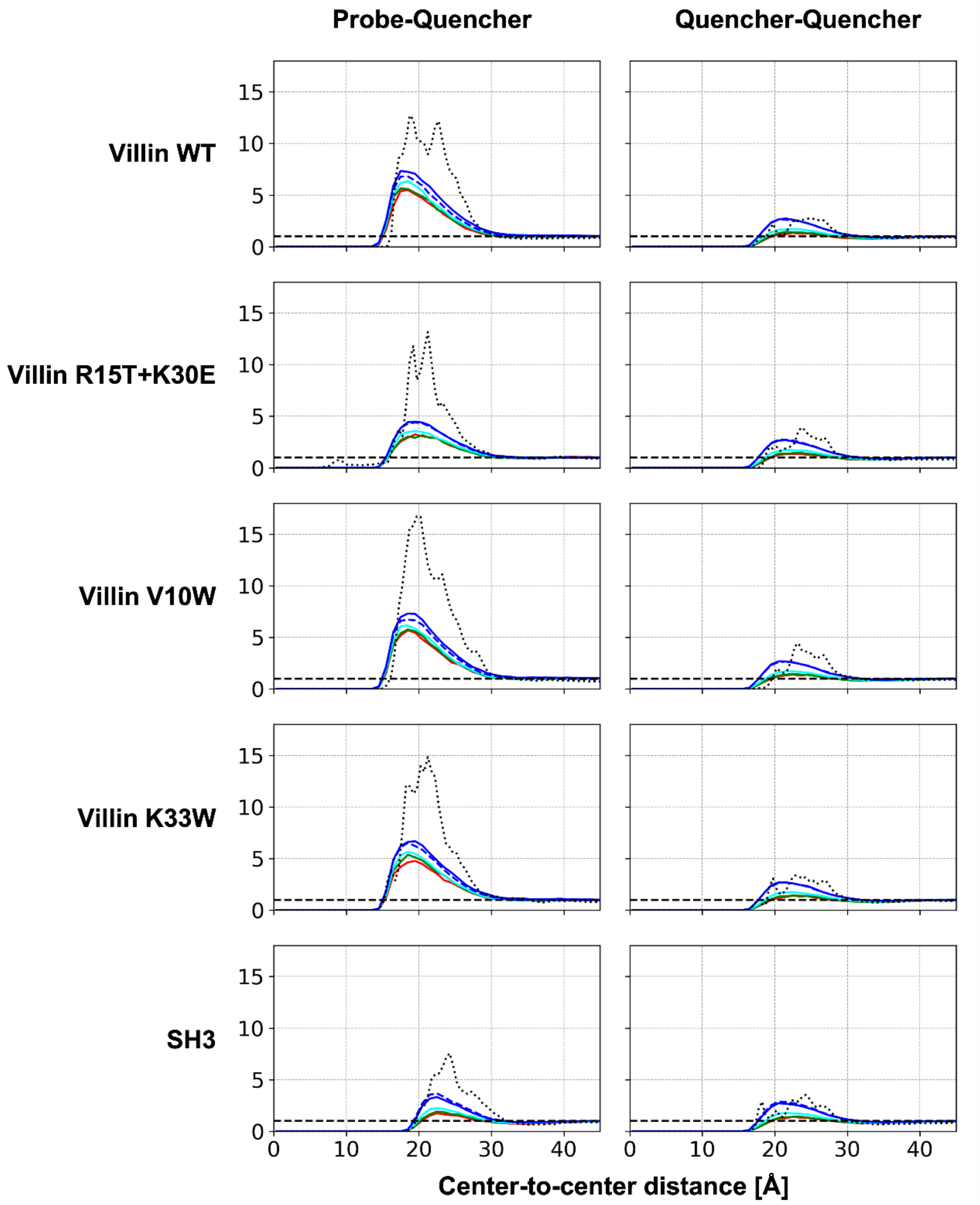


**Figure S15.** Radial distribution functions between proteins from the coarse-grained simulations. The quencher concentrations of 0.5, 0.8, 2.0, and 5.0 are shown in red, green, cyan, and blue, respectively. Data for the all-atom and coarse-grained simulations with the same composition (one probe and nine quencher proteins) are compared in black dotted and blue dashed lines, respectively. Black dashed lines are overlaid for g(r)=1.

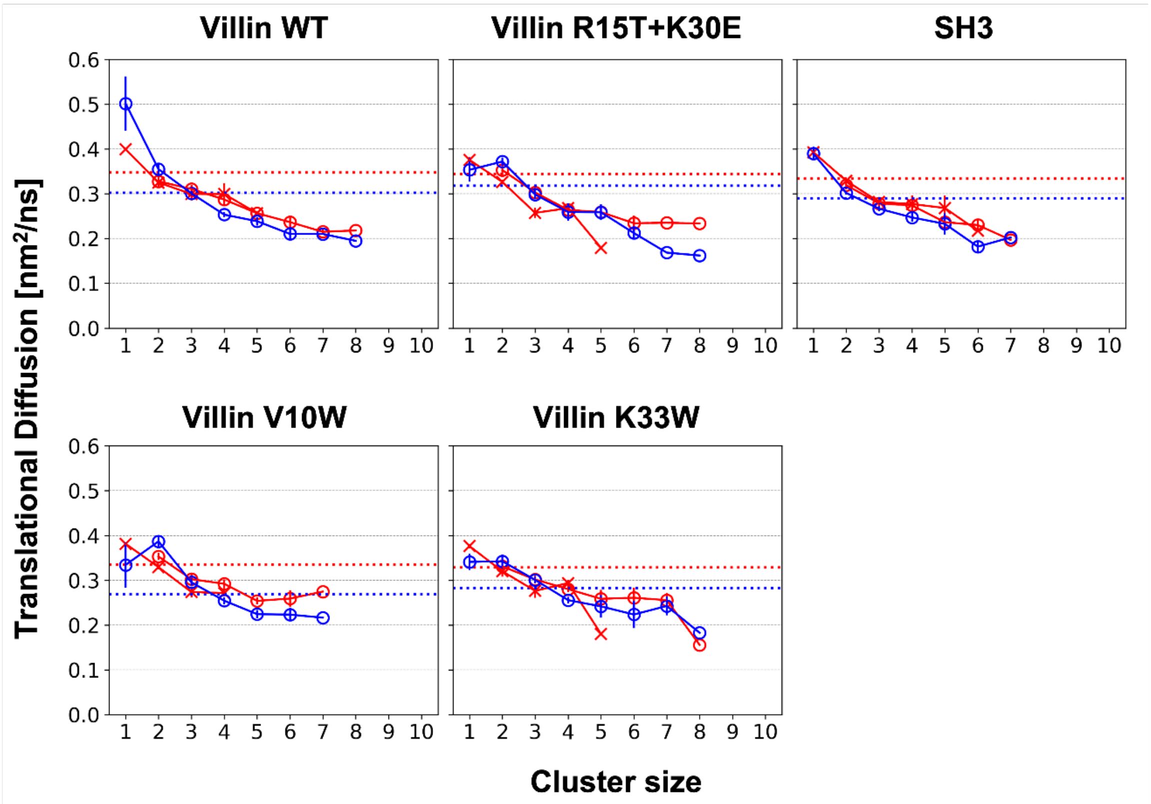


**Figure S16.** Translational diffusion coefficients as a function of protein cluster size. Cluster sizes that lasted shorter than 10 ns were discarded from the analysis. Data for probe and quencher proteins are shown in blue and red, respectively. For quencher protein, data for clusters with and without probe protein are marked with circles and Xs, respectively. Translational diffusion coefficients regardless of the cluster size are noted with horizontal dotted lines.

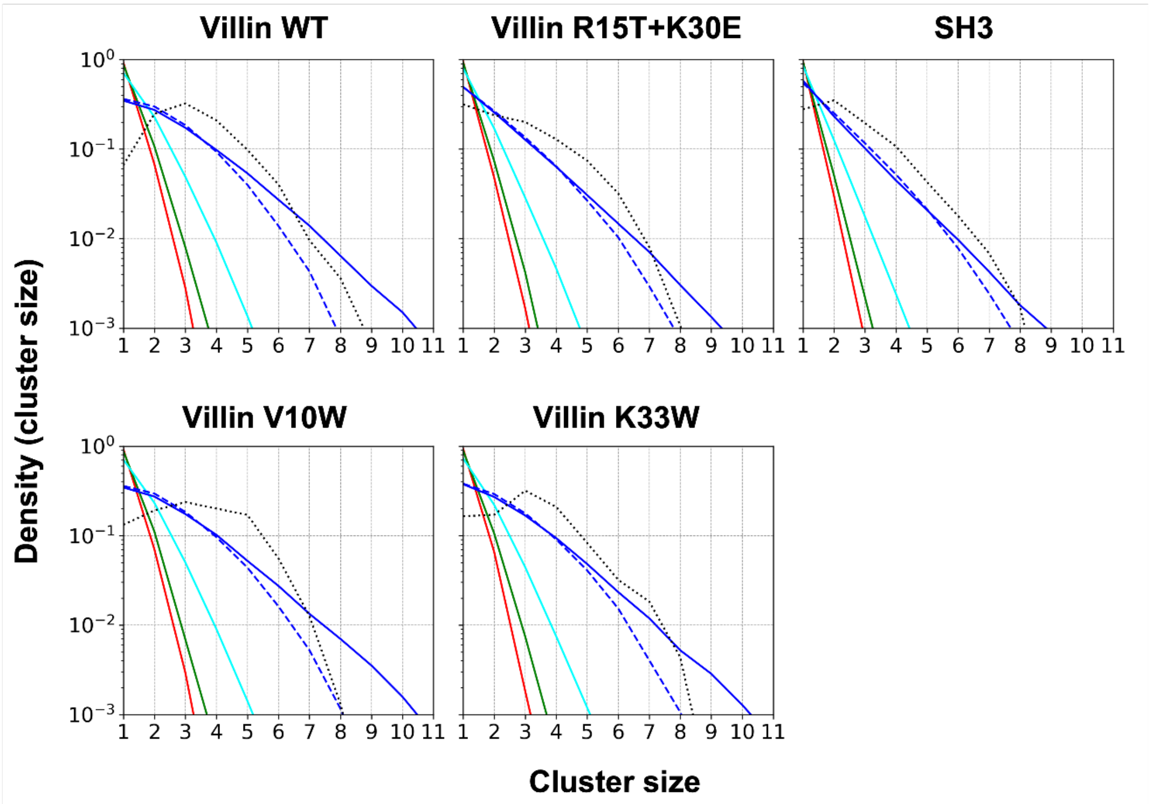


**Figure S17.** Cluster size distributions from CG simulations at different concentrations (solid lines colored as in **Fig. S15**) compared with atomistic simulation results in dotted lines. The dashed lines indicate results for CG simulations with a smaller number of molecules matching the atomistic simulations.

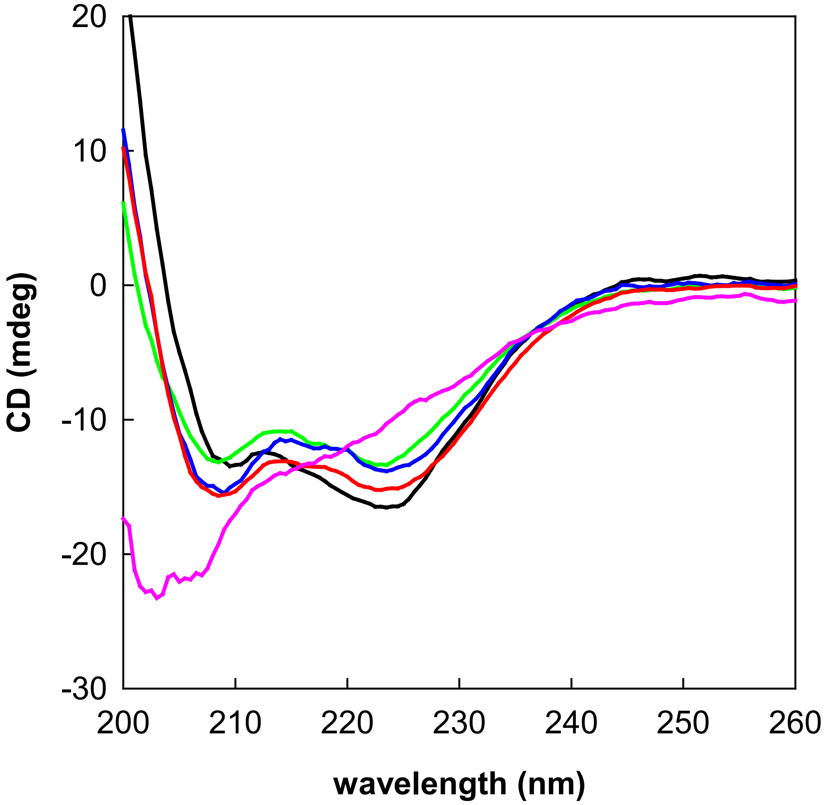


**Figure S18**. Circular dichroism spectra of villin headpiece HP35 wildtype (black), V10W (red), K33W (blue), R15T+K30E (green) and the SH3 domain (magenta). Amplitudes of the villin mutants were scaled to correct for concentration and normalized to the concentration of wildtype (15.5 μM).

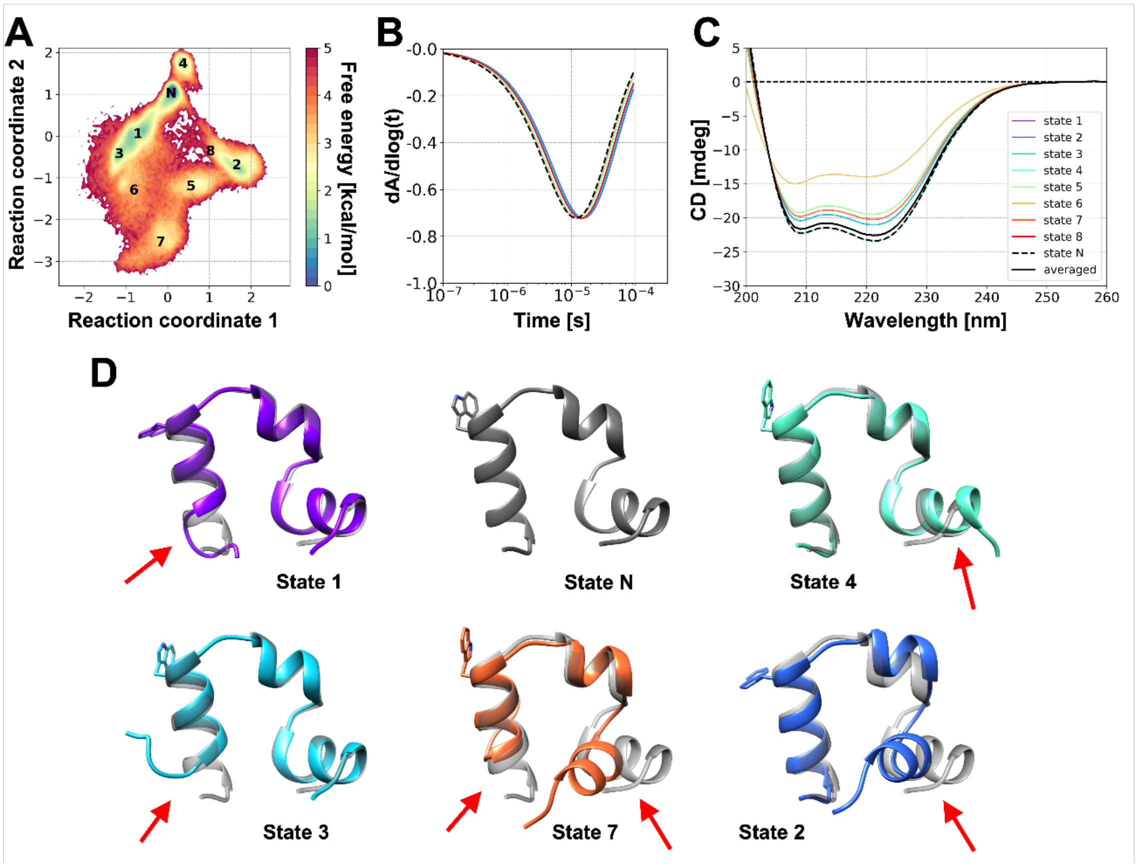


**Figure S19.** Diverse conformational states of villin wild-type and survival probability for the states. A: Free energy landscape of the protein and assigned conformational states from the Markov state modeling analysis. The native state is marked as "N", while the other states are noted with numbers. B: The slope of the survival probability curves for the probe protein in different conformational states. C: Predicted circular dichroism (CD) spectra for the states by SESCA (14) with a basis set of DSSP-1SC3. Data for the native state is shown as black dashed line (B, C), while state population-weighted averaged spectra is shown as black solid line (in C). D: Representative structures for several selected states are shown in the cartoon representation. The Tryptophan residue is additionally shown in stick representation. The native state conformation is overlaid in transparent gray on structures for the other states, and key structural differences are indicated by red arrows.

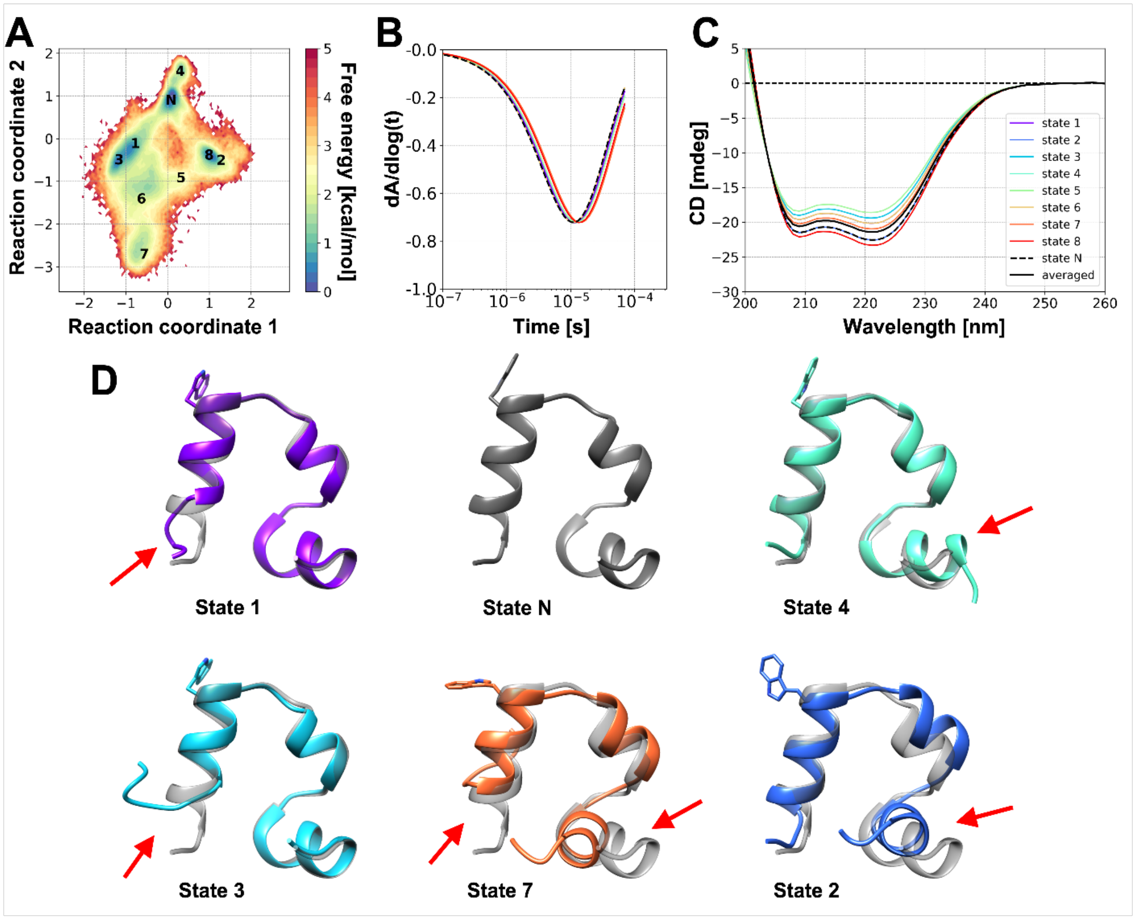


**Figure S20.** Free energy landscape and representative structures for conformational states of villin R15T+K30E. See **Fig. S19** for detailed descriptions.

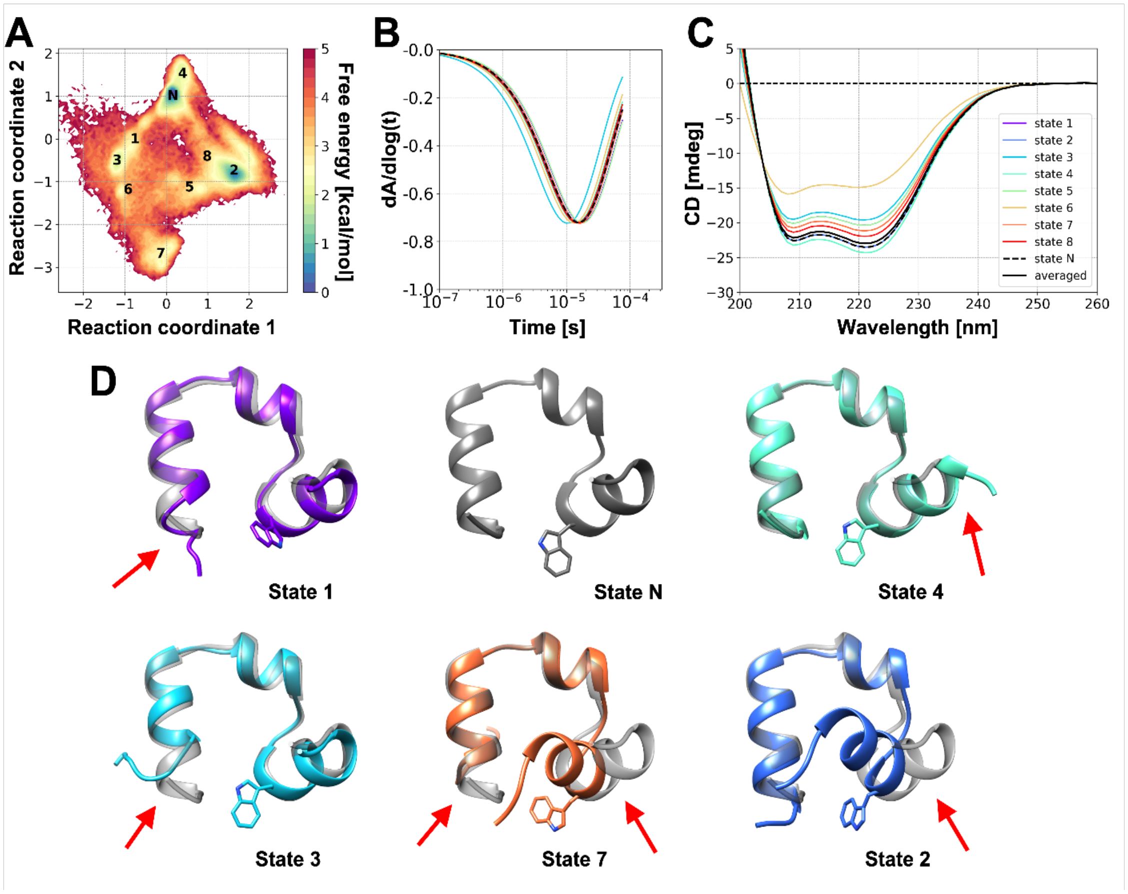


**Figure S21.** Free energy landscape and representative structures for conformational states of villin V10W. See **Fig. S19** for detailed descriptions.

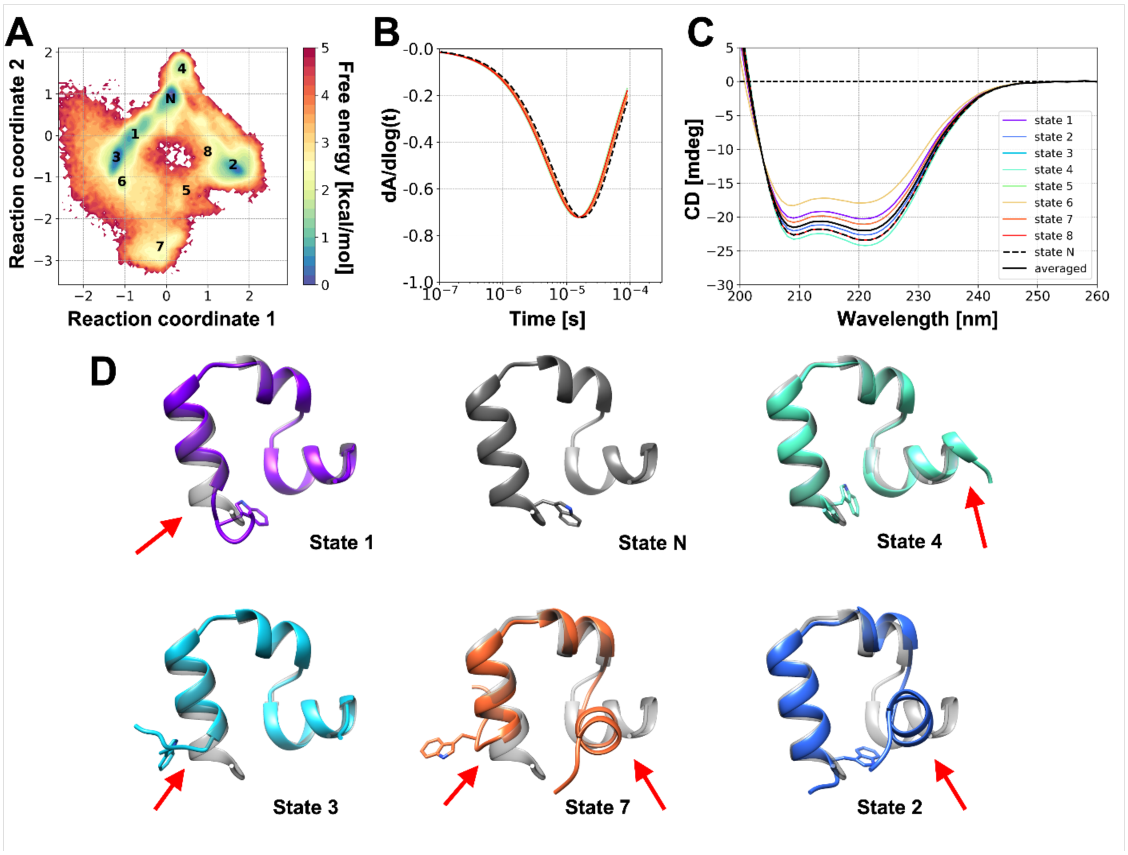


**Figure S22.** Free energy landscape and representative structures for conformational states of villin K33W. See **Fig. S19** for detailed descriptions.

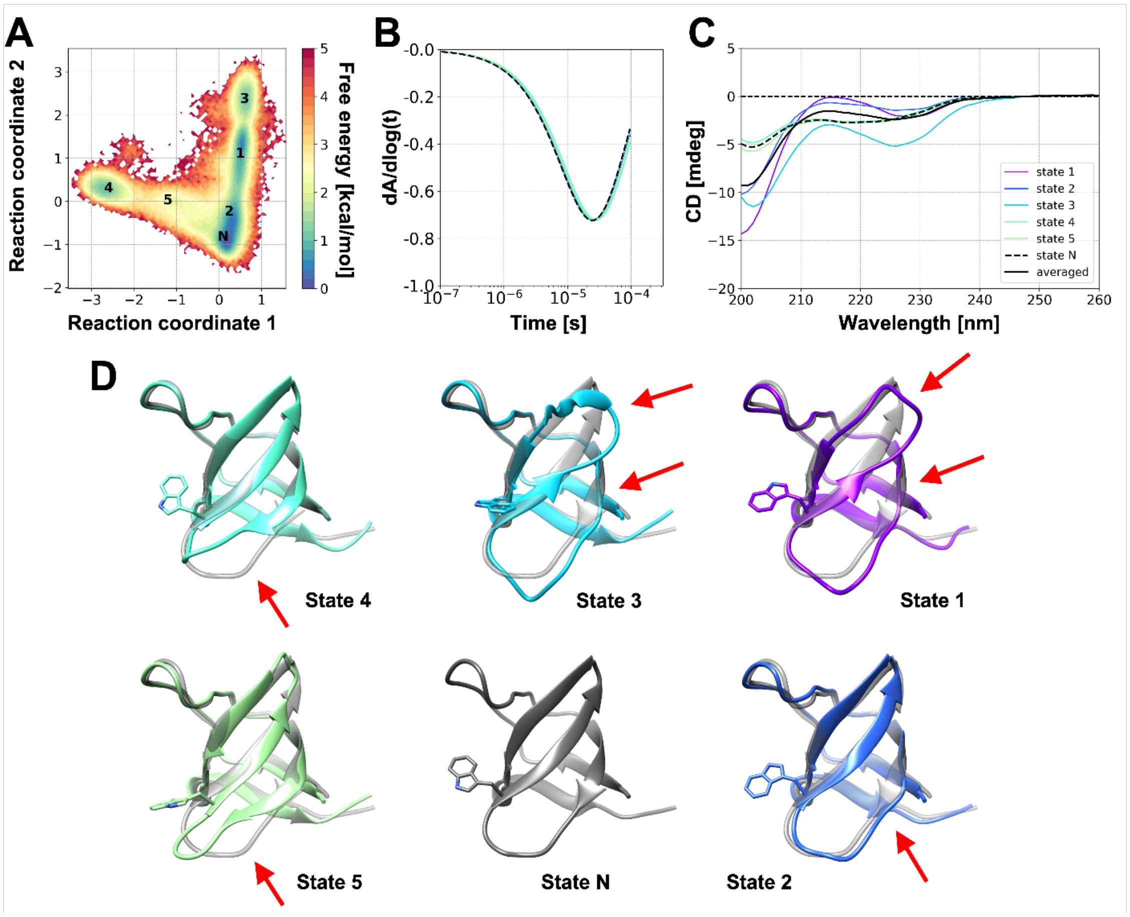


**Figure S23.** Free energy landscape and representative structures for conformational states of SH3. See **Fig. S19** for detailed descriptions.

**Table S1. Minima in derivative curves from experiments**

| Probe | dAbsorbance vs. log(t) Minimum | |
|---|---|---|
| | **Value** | **Time [μs]** |
| Villin WT | -0.493 ± 0.020 | 11.8 |
| Villin V10W | -0.620 ± 0.016 | 8.9 |
| Villin K33W | -0.650 ± 0.015 | 10.5 |
| Villin R15T+K30E | -0.554 ± 0.015 | 10.5 |
| SH3 | -0.534 ± .009 | 39.1 |

Values were extracted from data shown in **Fig. 2B** by fitting a fourth order polynomial functions. The errors are estimated from the derivative of the experimental data, obtained as a linear fit to the points around the minimum.

**Table S2. Translational diffusion coefficients for proteins and time-scale correction factors**

| System | Force Field | $D_{t,MD}$ [nm²/ns] | | $D_t$ [nm²/ns] | | $f_{TIP3P,PBC}$ (*Eq.* S2) |
|---|---|---|---|---|---|---|
| | | ***Probe*** | ***Quencher*** | ***Probe*** | ***Quencher*** | |
| **Villin WT** | **c36** | 0.164 (0.011) | 0.229 (0.005) | 0.273 (0.011) | 0.338 (0.005) | 1.726 |
| | **c36 + water** | 0.261 (0.012) | 0.285 (0.006) | 0.370 (0.012) | 0.393 (0.006) | 1.840 |
| | **c36m** | 0.193 (0.013) | 0.239 (0.004) | 0.302 (0.013) | 0.348 (0.004) | 1.749 |
| | **c36mw** | 0.274 (0.011) | 0.276 (0.004) | 0.356 (0.011) | 0.385 (0.004) | 1.825 |
| **Villin V10W** | **c36m** | 0.160 (0.008) | 0.226 (0.005) | 0.269 (0.008) | 0.334 (0.005) | 1.742 |
| **Villin K33W** | **c36m** | 0.174 (0.009) | 0.220 (0.004) | 0.283 (0.009) | 0.328 (0.004) | 1.718 |
| **Villin R15T+K30E** | **c36m** | 0.209 (0.018) | 0.236 (0.005) | 0.318 (0.018) | 0.344 (0.005) | 1.703 |
| **SH3** | **c36m** | 0.182 (0.010) | 0.226 (0.004) | 0.290 (0.010) | 0.334 (0.004) | 1.721 |

**Table S3. Amino acid sequences for proteins described in this work**

| Protein | Sequence |
|---|---|
| Villin WT (N28H) | MLSDEDFKAV FGMTRSAFAN LPLWKQQHLK KEKGLF |
| Villin V10W | MLSDEDFKAW FGMTRSAFAN LPLYKQQNLK KEKGLF |
| Villin K33W | MLSDEDFKAV FGMTRSAFAN LPLYKQQNLK KEWGLF |
| Villin R15T+K30E | MLSDEDFKAV FGMTTSAFAN LPLWKQQHLE KEKGLF |
| SH3 T22G | MEAIAKHDFS ATADDELSFR KGQILKILNM EDDSNWYRAE LDGKEGLIPS NYIEMKNHD |
| Protein G (K10C) | MTYKLILNGC TLKGETTTEA VDAATAEKVF KQYANDNGVD GEYTYDDATK TFTVTE |

The first methionine residue (highlighted in green) was not presented in the experimental constructs for villin. Tryptophan residues and point mutations are highlighted in blue and red, respectively.

**Table S4. MD simulation summary**

| System | Force Field | Simulation lengths |
|---|---|---|
| **Villin WT** | **c36** | Total: 36.6 μs<br>Anton: 3 x 3.0 μs = 9.0 μs<br>GPU: 5 x 3.6 μs + 4 x 2.4 μs = 27.6 μs |
| | **c36 + water** | Total: 9.0 μs<br>Anton: 3 x 3.0 μs = 9.0 μs |
| | **c36m** | Total: 42.6 μs<br>Anton: 1 x 6.0 μs + 3 x 3.0 μs = 15.0 μs<br>GPU: 1 x 3.6 μs + 8 x 3.0 μs = 27.6 μs |
| | **c36mw** | Total: 22.0 μs<br>Anton: 1 x 4.0 μs + 6 x 3.0 μs = 22.0 μs |
| **Villin V10W** | **c36m** | Total: 33.6 μs<br>Anton: 1 x 6.0 μs + 1 x 3.6 μs = 9.6 μs<br>GPU: 8 x 3.0 μs = 24.0 μs |
| **Villin K33W** | **c36m** | Total: 33.6 μs<br>Anton: 1 x 6.0 μs + 1 x 3.6 μs = 9.6 μs<br>GPU: 8 x 3.0 μs = 24.0 μs |
| **Villin R15T+K30E** | **c36m** | Total: 33.6 μs<br>Anton: 1 x 6.0 μs + 1 x 3.6 μs = 9.6 μs<br>GPU: 8 x 3.0 μs = 24.0 μs |
| **SH3** | **c36m** | Total: 36.0 μs<br>Anton: 2 x 6.0 μs = 12.0 μs<br>GPU: 8 x 3.0 μs = 24.0 μs |

**Table S5. MC parameters used to match atomistic simulation data**

| | **Villin WT** | **Villin V10W** | **Villin K33W** | **Villin R15T+K30E** | **SH3** |
|---|---|---|---|---|---|
| $\sigma$ [Å] | 3.28 | 3.25 | 3.42 | 3.40 | 3.85 |
| $\varepsilon$ [kcal/mol] | 6.30 | 4.80 | 1.30 | 2.95 | 5.00 |
| $\varepsilon_l$[1][kcal/mol] | 7.00 | 7.00 | 7.00 | 7.00 | 1.00 |
| potential | 12-6 | 12-6 | 12-6 | 12-6 | 10-5 |
| a [kcal/mol] | 0.75 | 0.45 | 1.20 | 1.20 | 1.20 |
| $\mu$ [Å] | 4.70 | 4.52 | 6.10 | 5.90 | 8.00 |
| w [Å] | 0.3 | 0.3 | 1.9 | 1.6 | 2.1 |
| $a_2$[2] [kcal/mol] | 0.3 | 0.9 | 0.5 | 0.5 | 0 |
| $\mu_2$[2] [Å] | 6.00 | 6.85 | 8.20 | 4.40 | - |
| $w_2$[2] [Å] | 0.35 | 1.40 | 0.40 | 0.30 | - |
| $D_{dh}$ [kcal/mol] | 25 | 11 | 17 | 20 | 0.4 |
| $K_{dh}$ [Å] | 3.5 | 4.0 | 4.0 | 4.0 | 60 |
| cutoff[3] [Å] | 10 | 9.5 | 10 | 9.5 | 10 |
| $v_a$[4] | 0.001 | 0.0035 | 0.0002 | -0.007 | 0.0001 |
| $v_b$[4] | 1.90 | 1.20 | 1.65 | 3.60 | 3.00 |

See **Eq. S1** for meaning of parameters.
[1]applied for distances smaller than contact minimum
[2]second Gaussian function
[3]distance at which Debye-Hückel potential becomes effective
[4]distance-scaled maximum step size: $\Delta x = 5$ Å$*(v_a*r)+v_b$

**Table S6. CD spectrum analysis from experiments**

| Probe | Helix | Sheet | Turn | Other |
|---|---|---|---|---|
| Villin WT | 64.5 | 0 | 16.3 | 19.2 |
| Villin R15T+K30E | 55.9 | 0 | 17.0 | 27.1 |
| Villin V10W | 56.4 | 0 | 20.5 | 23.1 |
| Villin K33W | 58.6 | 0 | 18.7 | 22.7 |
| SH3 | 15.7 | 22.7 | 16.7 | 44.9 |

**Table S7. Secondary structure analysis from simulations**

| Probe | Helix | Sheet | Coil |
|---|---|---|---|
| Villin WT | 62.1 *(5.0)* | 0.1 *(0.6)* | 37.8 *(4.9)* |
| Villin R15T+K30E | 59.9 *(5.2)* | 0.0 *(0.2)* | 40.1 *(5.2)* |
| Villin V10W | 60.2 *(6.1)* | 0.1 *(0.6)* | 39.7 *(5.9)* |
| Villin K33W | 60.1 *(5.7)* | 0.0 *(0.3)* | 39.9 *(5.7)* |
| SH3 | 3.8 *(2.7)* | 42.1 *(3.3)* | 54.1 *(4.3)* |

Average secondary structure contents in % based on atomistic simulation snapshots analyzed with DSSP (15). Standard deviations are given in parentheses.
